# A Tiered Approach to Human Synapse Proteomics: Optimized LC-MS/MS Analysis of Whole-Tissue and Synaptosome Preparations from Frozen Post-Mortem Brain Samples

**DOI:** 10.1101/2025.06.04.657790

**Authors:** Femke C. Roig-Kuhn, Remco V. Klaassen, Frank T.W. Koopmans, Tiara S.Z. Koolman, August B. Smit, Sabine Spijker

## Abstract

Recent advancements in neuroproteomics have enabled detailed analysis of protein expression and function in the human brain. Post-mortem human brain studies have significantly advanced our understanding of the relationship between genetics, cell biology of neurological and psychiatric disorders and their clinical diagnosis. Synaptic dysfunction often has a central role in these disorders. Therefore, we specifically evaluated the sensitivity of liquid chromatography-tandem mass spectrometry (LC-MS/MS) to detect synaptic proteins in whole-tissue lysates versus synaptosome preparations. First, we optimized sample preparation protocols for frozen human gray matter (GM), refining the suspension TRAPping (sTRAP) digestion method to improve protein solubilization and Cysteine reduction and alkylation using thin human tissue sections, and to accomplish low technical variation by minimizing sample handling. We achieved a highly reproducible sample preparation workflow by rigorously applying standardization and randomization across dissection, processing, and LC-MS/MS runs. Second, comparative LC-MS/MS analysis showed that cortical whole-tissue lysates are a practical solution for large-scale studies and broadly detected synaptic proteins focusing on excitatory neurons. However, enrichment by synaptosome isolation offered improved resolution of synapse-specific proteins. Because synapse-proteomics enables insight into spatial regulation—i.e., alterations at the synapse that are not reflected in the soma–we recommend a tiered approach: initial whole-tissue analysis for broad disease-associated changes, followed by targeted synaptosome proteomics to deepen insight into synaptic alterations. This strategy optimally balances throughput, reproducibility, and biological relevance, and enhances the study of brain disorders through proteomics. Moreover, analyzing synaptic proteins first at the tissue level improves insight into overall regulation of synaptic proteins induced by synapse loss or gain.

## 1. Introduction

Molecular techniques to study neurodegenerative and neuropsychiatric disorders are becoming increasingly precise and comprehensive. With the advent of high-throughput omics analyses like transcriptomics and proteomics, it is now possible to characterize the molecular signatures that distinguish specific disorders from healthy controls in large patient cohorts. This aids tremendously in uncovering underlying disease mechanisms, as a prelude to researching new treatment options. Yet, quantitative analyses on large cohorts pose many questions on how to best prepare and analyze the samples in a manner tailored to the specific research project.

One of the most direct methods to investigate differences between health and disease in humans is using frozen post-mortem brain tissue, made available by brain banks worldwide, with the goal of increasing the understanding of the human brain [1]. In addition to limitations in the quantity of brain tissue available from brain banks, omics studies face several key challenges. Chief among these are issues related to tissue quality—especially the post-mortem interval—which can lead to degradation of RNA and proteins. Sample heterogeneity, including differences in age, sex, cause of death, medication use, and comorbid conditions, further complicates data interpretation. Technical challenges in sample preparation, such as variability and batch effects, also pose significant obstacles. These factors are especially critical in research on neuropsychiatric disorders, which often lack clear neuropathological markers. Moreover, proteomic alterations in these conditions are typically subtle and highly variable, reflecting the inherent biological complexity of human brain tissue. Careful experimental design, rigorous methodological controls, and minimizing technical variation — particularly during protein extraction and mass spectrometry [2] — are therefore essential to obtain meaningful insights.

Proteomics protocols are widely established in rodent research [3,4]. However, mouse tissue differs from human tissue in several important anatomical and biochemical aspects that should be considered when translating these mouse-protocols to human research. Compared to mice, humans exhibit slightly higher neuronal density, increased cortical thickness and larger neurons [5], along with a substantially greater brain volume driven by a more expanded and folded cerebral cortex. Additionally, human brain tissue contains a higher myelin content than mouse brain tissue [6]. Because myelin is also present in human grey matter [7], the altered protein-lipid composition of human brain tissue necessitates protocol modifications, such as sonication [8], to ensure efficient protein extraction. However, introducing additional steps could increase technical variability between batches. To mitigate this, we sought to adapt existing protocols to improve sample solubilization and preparation for mass spectrometry, while minimizing additional steps that would compromise reproducibility.

In addition to whole-tissue lysate proteomics, measurements on subcellular fractions –coined under the umbrella of spatial proteomics [9]– provides valuable insights into neuronal function and pathology. In the brain, synaptic connections are underlying neural network physiology, and synaptic transmission can be affected under disease conditions. Synaptic alterations have been associated with neurodegenerative disorders [10,11] and neuropsychiatric disorders [12–14]. As such, synaptic fractionation is a powerful approach to study disease-specific regulation at the synapse. Isolating the synaptic compartment allows detection of subtle changes in proteins specific to the synapse that would otherwise not be detected in the lysate due to dilution in the whole-tissue context. Historically, synaptic fractionation reduces sample complexity, making it possible to detect more low-abundant synaptic proteins with mass spectrometry. Over the past 6 years, several human proteomics studies focusing on the synapse have been published using different biochemical fractionation approaches (Table S1, [15–18]), with several studies detecting 5,000-9,000 proteins. These studies make it important to evaluate the extent of enrichment of synaptic proteins in synaptic isolations relative to whole-tissue preparations. Previous research in mice has revealed that brain region-specific differences in synaptic protein enrichment [19]. The extent to which these differences translate to human brain tissue is unclear, as currently there are no studies that compare the whole cell lysate proteome to that of the synaptosome with a detection yield of >5,000 proteins.

To evaluate the sensitivity of liquid chromatography-tandem mass spectrometry (LC-MS/MS) for detecting synaptic proteins in human whole-tissue lysates versus synaptosome preparations, we first implemented a standardized and randomized workflow designed to minimize sample handling and reduce batch effects. Limiting sample handling enhances reproducibility and increases the feasibility of large-scale studies with several hundreds of samples [20–22]. Here, we adapted a revised version of the original ProtiFi suspension-trapping (S-TRAP) mini spin column digestion protocol 4.1 previously described by our lab [3]. This revised ‘sTRAP’ version replaced the commercial Protifi S-TRAP column with a low-cost plasmid DNA micro-spin column, which demonstrated superior performance in number of identified proteins and peptides, as well as lower coefficients of variation (CoV). With the aim of optimizing reproducible proteomic profiling of human brain tissue, we systematically evaluated and adapted the existing sTRAP workflow to processing micrometer-thin tissue sections in a one-tube reaction. We tested the effect of incubation conditions and centrifugation on extract composition and explored the impact of variable tissue input levels on tryptic digestion efficiency and the depth and consistency of protein-level quantification. Throughout the workflow, the sequence of dissection, sample preparation, and LC-MS/MS analysis were carefully arranged and randomized to avoid potential confounding factors. Finally, we explored whether isolating the synaptic fraction provides added value in the detection of low-abundant synaptic proteins in a set of control donors.

Together, we describe 1) an optimized digestion protocol to maximize peptide and protein yield for LC-MS/MS using thin human post-mortem cryo-sections, and 2) a comparison between whole-tissue and synaptosome proteomics, exploring the added advantages of enriching for synapses given recent improvements in mass spectrometry sensitivity. By outlining the rationale behind each step, we aim to support researchers in designing and executing robust and reproducible quantitative proteomic analyses on post-mortem brain tissue.

## 2. Materials and Methods

### 2.1. Tissue Collection from Human Brain Samples

Tissue was obtained from the Netherlands Brain Bank (NBB). Informed consent was obtained from all donors and their next of kin for the use of material and clinical data for research purposes, as outlined in https://www.brainbank.nl/about-us/ethics/. The NBB autopsy procedures and the use of tissue for research were approved by the Ethics Committee of Amsterdam UMC (registered with the US Office of Human Resource Protections as IRB00002991 under Federal wide Assurance number 00003703), location VUmc, Amsterdam, The Netherlands, albeit that according to the Dutch Act on Medical Research Involving Human Subjects (WMO) no ethical approval is required for collection nor use of donated brain tissue.

For optimizing the proteomics workflow, we used tissue from the most anterior part of the superior frontal gyrus (SFG) –comprising sections 2-7, as presented in the Adult Human Allen Brain Reference Atlas [23], overlapping with the medial and lateral subdivision of Brodmann area 9 and 10– of two replicate tissue samples from a non-demented control donor (NDC), and two replicate tissue samples from a donor diagnosed with (presenile) Alzheimer’s Disease (AD) with (putative) comorbid schizophrenia (Supplemental file 1, clinical information sheet). For analysis of whole-tissue lysates vs. synaptosome preparations, we used the most anterior part of the SFG from 8 NDC donors (Table 1). For all samples, gray matter (GM) was collected from tissue sections (2–10 mg cut at 10 µm for whole-tissue lysate; ∼25 mg cut at 50 µm for synaptosome preparations) using a cryostat.For optimizing the proteomics workflow, we used tissue from the most anterior part of the superior frontal gyrus (SFG) –comprising sections 2-7, as presented in the Adult Human Allen Brain Reference Atlas [23], overlapping with the medial and lateral subdivision of Brodmann area 9 and 10– of two replicate tissue samples from a non-demented control donor (NDC), and two replicate tissue samples from a donor diagnosed with (presenile) Alzheimer’s Disease (AD) with (putative) comorbid schizophrenia (Supplemental file 1, clinical information sheet). For analysis of whole-tissue lysates vs. synaptosome preparations, we used the most anterior part of the SFG from 8 NDC donors (Table 1). For all samples, gray matter (GM) was collected from tissue sections (2–10 mg cut at 10 µm for whole-tissue lysate; ∼25 mg cut at 50 µm for synaptosome preparations) using a cryostat.

**Table 1.**
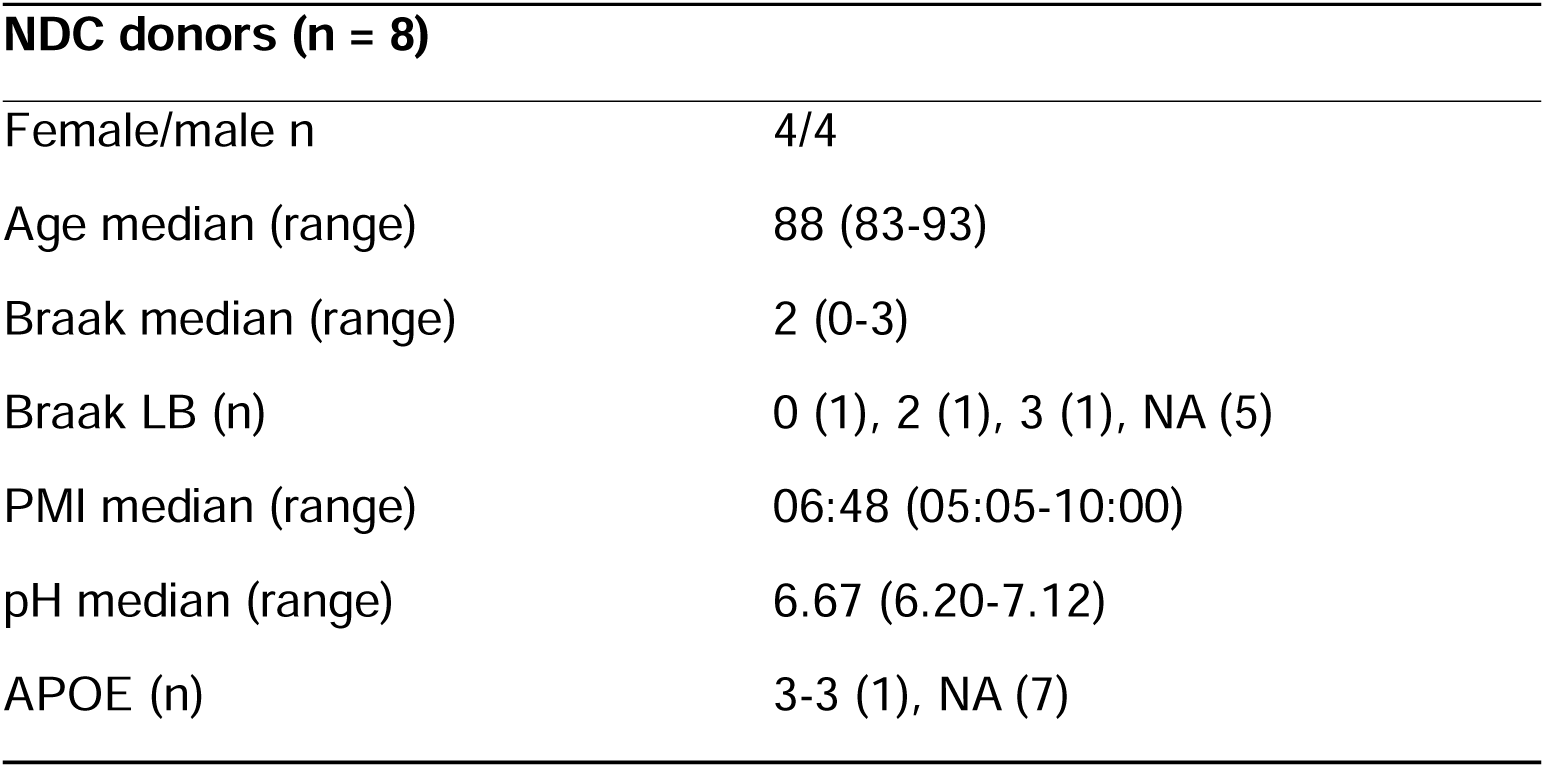
Demographic and clinical information of the superior frontal gyrus samples used in subcellular fraction isolation (synaptosome proteomics). NDC: non-demented control. NA = not measured. Braak LB = Braak Lewy Bodies. PMI = post-mortem interval. APOE = apolipoprotein E allele.

| **NDC donors (n = 8)** |  |
| --- | --- |
| Female/male n | 4/4 |
| Age median (range) | 88 (83-93) |
| Braak median (range) | 2 (0-3) |
| Braak LB (n) | 0 (1), 2 (1), 3 (1), NA (5) |
| PMI median (range) | 06:48 (05:05-10:00) |
| pH median (range) | 6.67 (6.20-7.12) |
| APOE (n) | 3-3 (1), NA (7) |

### 2.2. Subcellular Fraction Isolation: Synaptosome Proteomics

For synaptosome isolation, a similar protocol was used as previously published by Pandya et al. 2017 [8] and [24], which is based on the protocol of [25] with minor modifications. In short: cortical GM tissue (∼25 mg, sectioned at 50 µm) was homogenized in 2 mL ice-cold homogenization buffer (0.32 M Sucrose, 5 mM HEPES pH 7.4) using a Teflon/glass homogenizer (Schuett homgenplus; Schuett-biotec 9651560) set at 900 rpm for 12 strokes. For a detailed and step-by-step description, see Supplemental protocol 2. Note that multiple considerations could be taken along for every step during synaptosome isolation; see [26]. The resulting homogenate was collected and centrifuged at 1,000x g for 10 minutes (4 °C) to spin down the nuclear membranes. The supernatant (S1) was placed on a discontinuous sucrose gradient consisting of 0.85 and 1.2 M sucrose. After ultracentrifugation at 187,000x g at 4 °C for 2 h, the interphase disk between the sucrose concentrations – containing the synaptosome fraction– was collected. After centrifugation (18,000x g (Eppendorf 5810) at 4 °C for 30 minutes), the pellet was resuspended in homogenization buffer, and the samples were set aside for sTRAP.

### 2.3. MS sample preparation using suspension trapping (sTRAP)

MS sample preparation was performed following the DNA micro spin column suspension trapping (sTRAP) protocol as described by our lab previously [3], with several adaptations to accommodate for human tissue samples.

Human brain sections – ∼10 mg for the whole-tissue extraction optimization, and 2-5 mg for comparison to the synaptosome fraction – were extracted in 500 µL sTRAP lysis buffer (5% SDS, 50 mM Tris-HCl (pH 8.0), 5 mM tris(2-carboxyethyl)phosphine (TCEP) and 20 mM 2-Chloroacetamide (2-CAA)) by incubation in a ThermoTop-covered Thermomixer set to 1700 rpm, either at 55 °C for 30 minutes or at 95 °C for 15 minutes followed by 55 °C for 15 minutes. This one-pot extraction procedure not only homogenized the thin tissue sections but also immediately reduced disulfide bonds and alkylated the free sulfhydryl residues for downstream MS-analysis. Insoluble debris was cleared from the tissue lysates by centrifugation at 20,000x g for 10 minutes at RT. For lysates belonging to the whole-tissue extraction optimization experiment, aliquots were collected both before and after centrifugation, providing the ‘PRE-centrifugation’ and ‘POST-centrifugation’ fractions, respectively. Synaptosome samples were processed using the same procedure, with a final extraction volume of 70 µL and a similar extraction condition of 95 °C for 15 minutes followed by 55 °C for 15 minutes. The protein concentration of lysates was determined using the bicinchoninic acid (BCA) protein assay.

For the sTRAP workflow, either 50 µg (four replicates for each sample for whole-tissue extraction optimization) or 10 µg (synaptosome proteomics for each donor) of total protein was used as input (in 50 µL sTRAP lysis buffer). For the Trypsin/Lys-C digestion efficiency test, pooled whole-tissue lysates were serially diluted to provide an sTRAP protein input of 50 µg, 25 µg, 12.5 µg and 5 µg. Samples were acidified to a final concentration of 1.1% phosphoric acid (12% stock solution), mixed with six volumes of binding/washing buffer (90% methanol in 100 mM Tris-HCl pH 8.0), and loaded onto a plasmid DNA micro column (HiPure from Magen Biotechnology - C13011, see [3]). The protein particulate was retained on the column upon centrifugation at 1,400x *g* for 1 minute and the columns were washed four times with binding/washing buffer. Columns were transferred to new LoBind tubes (Eppendorf), supplemented with either 1 µg Trypsin/Lys-C (whole-tissue extraction optimization; Promega - V507A) or 0.4 µg Trypsin (synaptosome proteomics) in 50 mM NH_4_HCO_3_ and incubated overnight at 37 °C in a humidified incubator. Tryptic peptides were eluted and pooled by subsequent addition of 50 mM NH_4_HCO_3_, 0.1% formic acid and 0.1% formic acid in acetonitrile. Collected peptides were dried by SpeedVac and stored at -80 °C.

For a detailed and step-by-step description of the optimized protocol, see Supplemental protocol 1.

### 2.4. LC-MS Analysis

Each sample of tryptic digest was redissolved in 0.1% formic acid, and the peptide concentration was determined by tryptophan-fluorescence assay [27]. Peptides were loaded onto an Evotip Pure (Evosep) according to the manufacturer’s instructions; 75 ng for the whole-tissue extraction optimization samples and 150 ng for the synaptosome proteomics samples. Peptides were separated by standardized 30 samples per day method on the Evosep One liquid chromatography system, using a 15 cm × 150 μm reverse-phase column packed with 1.5 µm C_18_-beads (EV1137 from Evosep) connected to a 20 µm ID ZDV emitter (Bruker Daltonics).

Peptides were electro-sprayed into the timsTOF Pro 2 mass spectrometer (whole-tissue extraction optimization; Bruker Daltonics), or the timsTOF HT mass spectrometer (synaptosome proteomics; Bruker Daltonics) equipped with CaptiveSpray source and measured with the following settings: Scan range 100-1700 m/z, ion mobility 0.65 to 1.5□Vs/cm^2^, ramp time 100 ms, accumulation time 100 ms, and collision energy decreasing linearly with inverse ion mobility from 59 eV at 1.6 Vs/cm^2^ to 20 eV at 0.6 Vs/cm^2^. Operating in dia-PASEF mode, each cycle took 1.38 s and consisted of 1 MS1 full scan and 12 dia-PASEF scans. Each dia-PASEF scan contained two isolation windows, in total covering 300-1200 m/z and ion mobility 0.65 to 1.50□Vs/cm^2^. Dia-PASEF window placement was optimized using the py-diAID tool [28]. Ion mobility was auto calibrated at the start of each sample (calibrant m/z, 1/K0: 622.029, 0.992 Vs/cm^2^; 922.010, 1.199 Vs/cm^2^; 1221.991, 1.393 Vs/cm^2^). Samples were processed in a randomized manner to prevent batch effects by MS run order (Supplemental Figure S1).

### 2.5. LC-MS Data Analysis

The dia-PASEF raw data were processed with DIA-NN (version 1.8.1; [29]). An in-silico spectral library was generated from the Uniprot human reference proteome (SwissProt and TrEMBL, canonical and additional isoforms, release 2023-02 (whole-tissue-extration optimization) and 2024-02 (synaptosome proteomics)) using Trypsin/P digestion and at most 1 missed cleavage. Fixed modification was set to carbamidomethylation (C), and variable modifications were oxidation (M) and N-term M excision (at most 1 per peptide). Peptide length was set to 7-30, precursor charge range was set to 2-4, and precursor m/z range was limited to 280–1220. Both MS1 and MS2 mass accuracy were set to 15 ppm, scan window was fixed at 9, and the precursor False Discovery Rate (FDR) was set to 1%. Heuristic protein inference was disabled, protein identifiers (isoforms) were used for protein inference, and double-pass mode was enabled. RT-dependent cross-run normalization was enabled with the robust LC quantification strategy. Match-between-runs (MBR) was enabled, but only between samples of the same fraction (i.e. whole-lysate or synaptosomes). All other settings were left as default.

MS-DAP (version 1.2.2; [30]) was used for downstream analysis. Peptide-level filtering was configured to retain only peptides that were confidently identified in at least 75% of the samples per group. Peptide abundance values were normalized using the VSN algorithm, followed by protein-level mode-between normalization. This approach was shown to be robust across a wide range of datasets, including those with strongly asymmetric protein abundance differences between experimental conditions [30]. Differential expression analysis was performed in a between-subject manner by the DEqMS algorithm [31]. The log_2_ foldchange (FC) threshold was estimated by bootstrap analysis for the optimization part (Supplemental file 1, Bootstrapping), as well as for the quantitative comparison of synaptosome vs lysate (-0.208 > log2FC > 0.208) and resulting P-values were adjusted for multiple testing using the Benjamini–Hochberg FDR procedure (FDR adjusted P-value cutoff of 1%). The effect size of a regulated protein represents the ratio of relative (log_2_) fold change to the mean abundance variance across replicates. Differential expression analysis for synaptosome vs lysate samples was performed accounting for relevant covariates (sex, age; see Table 1). Gene set enrichment analysis was done using the online tools ShinyGO (v0.85.1 [32]) and SynGO (v1.3) [33], always using the complete list of confidently identified proteins as background. For SynGO, we focused on gene ontology terms for cellular components in the synapse. ShinyGO was used with standard settings (FDR cutoff 5%) except for increasing the minimal pathway size from 2 to 3 proteins. Venn diagrams were made with InteractiVenn [34].

## 3. Results

In this study, we employed tissue sectioning to selectively collect GM from frozen human brain samples. This method is relatively straightforward and allows for precise anatomical targeting. Moreover, the use of thin (10 µm) tissue sections results in a high surface area-to-volume ratio, which facilitates efficient chemical solubilization. We first optimized our pre-MS workflow on thin GM sections using a selected set of donors (n=4) and subsequently applied this protocol to compare whole-tissue lysates with synaptosome-enriched preparations of NDC donors (n=8) (Figure 1).

**Figure 1.**
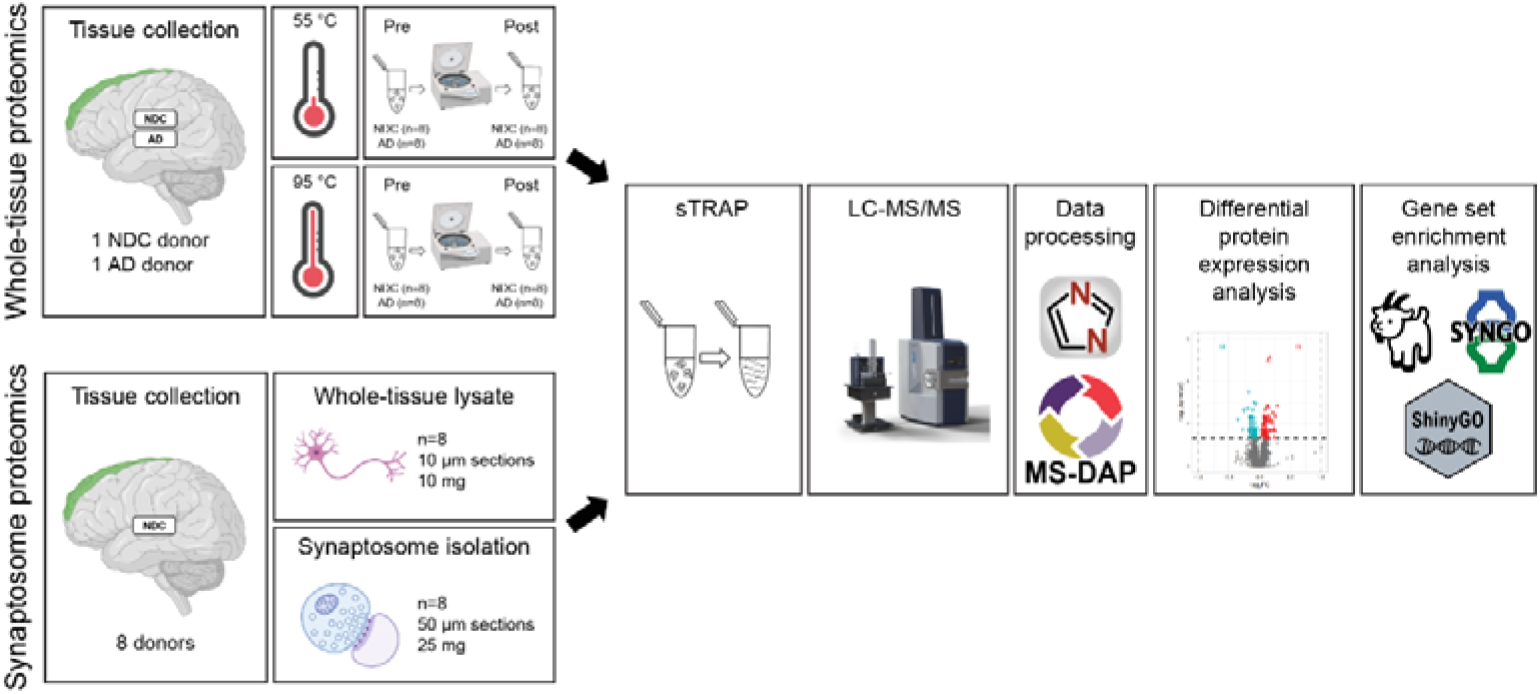
Overview of the proteomics analyses. For whole-tissue extraction optimization proteomics (above), two samples were obtained from each donor (10 µm sections, 10 mg per sample). Tissue extraction was performed at 55 °C and 95 °C (n=4 per condition; 2 NDC and 2 AD serial tissue samples). Protein samples were collected before and after centrifugation, yielding n=8 replicates per condition. Samples were then prepared for mass spectrometry with sTRAP, and the data was processed and analyzed. For the subcellular fraction isolation (below), two samples were collected per NDC donor: one for whole-tissue lysate proteomics (n=8, 10 µm sections, 10 mg per sample) and one for synaptosome proteomics (n=8, 50 µm sections, 25 mg per sample). Samples were then prepared for mass spectrometry with sTRAP, and the data was processed and analyzed. NDC = non-demented control donor; AD = donor with Alzheimer’s Disease and comorbid schizophrenia; Pre = pre-centrifugation; Post = post-centrifugation; sTRAP = suspension TRAPping; LC-MS/MS = liquid chromatography-tandem mass spectrometry.

### 3.1. Optimization of the sTRAP Micro Spin Column Digestion Protocol for LC-MS/MS Proteomics Analysis of Sectioned Post-Mortem Tissue

To perform neuroproteomic analyses of fresh-frozen human GM cortical tissue using a streamlined one-tube workflow with minimal sample handling, we adapted the recently published sTRAP micro spin column digestion protocol [3] to our sample input type (human GM) and quantity. We first optimized the lysis temperature and SDS buffer volume, changed the alkylation reagent [35], and tested Trypsin / Lys-C digestion conditions to ensure optimal protein solubilization and digestion. Protein solubilization is critical for ensuring replicability of samples, as variation in input material largely impacts experimental outcomes. This is specifically important in neurodegenerative research, where protein aggregates (e.g. plagues, tangle) are frequently observed and often constitute key pathological features for molecular analysis [36]. To account for these factors in the optimization part of our study, we used protein samples derived from 10 mg tissue to avoid exceeding solubilization limits. Moreover, tissue was collected from an NDC and AD donor, anticipating that AD samples might present a higher degree of protein aggregation, potentially affecting solubilization efficiency.

We first tested qualitative aspects of the solubilization protocol, focusing on SDS buffer volume and solubilization temperature. We avoided adding extra mechanical solubilization methods that could introduce sources of sample variation. Instead, we aimed at improving the chemical solubilization in SDS buffer by making the collected tissue sections thinner (10 µm) thereby increasing the surface area in contact with the buffer. Furthermore, we used a relatively high SDS buffer volume (500 µL), as this was sufficient to solubilize all the tissue without reaching the 1.5 mL Eppendorf tube volume limit. We experienced that lower volumes resulted in incomplete dissolution into the SDS buffer. The optimal temperature for solubilization was 95 °C for all sample types (NDC and AD). However, this caused an undesired pressure build-up during heating. To mitigate this, the temperature was set to 95 °C for 15 minutes, followed by 55 °C for another 15 minutes, maintaining the original 30-minute solubilization time.

Next, we quantified protein yield from a 10 mg tissue sample to determine the maximum amount of protein for digestion and calculated the amount of Trypsin / Lys-C accordingly to represent the lowest possible enzyme-protein ratio of 1:50. A 10 mg tissue sample represented the maximum tissue weight acquired during tissue collection, with average samples ranging between 2–5 mg. BCA assays showed no substantial difference in protein content between NDC and AD samples (Supplemental Figure S2) with approximately 9% of the tissue weight corresponding to protein (± 0.9 mg protein per 10 mg GM tissue). Based on this, a fixed quantity of 1 µg Trypsin / Lys-C was applied to all samples, maintaining an enzyme-to-protein ratio within the commonly used range of 1:10 to 1:25 (w/w) [3].

Subsequently, we tested the effectiveness of the 95 °C solubilization step during protein extraction compared to 55 °C in terms of protein yield, as measured quantitatively by MS analysis. Furthermore, we evaluated whether centrifugation after extraction was beneficial to obtain a more representative sample from a larger protein pool, as this step could remove larger insoluble aggregates and make the sample more homogeneous. On the other hand, it could result in the loss of specific proteins. For these tests, we compared technical replicates of the previous NDC and AD donors that were either centrifuged or not after solubilization at 55 °C or 95 °C (2 temperature settings, each with n=2 tissue replicates, each with n=4 sample replicates (PRE-centrifugation, POST-centrifugation), yielding 8 technical replicates per donor per condition).

Solubilization at 95 °C yielded approximately the same protein and peptide counts compared to 55 °C in NDC and AD, along with more consistent results in yield (Supplemental Figure S3). Added benefits of solubilization at 95 °C were the increased detection of peptides with modified Cysteine residues (11.0–11.9% at 55 °C to 15.9% at 95 °C; Table S2), indicating improved reduction-alkylation efficiency, and the slight decrease in coefficient of variation, apparently reducing (technical) variability between samples (Figure 2A). Higher temperature solubilization led to increased abundance levels (enriched at 95 °C) for a substantial portion of the detected proteins (NDC: 469/5741 = 8.64%; AD: 244/5701 = 4.28%), while only a small fraction showed decreased abundance levels (enriched at 55 °C; NDC: 26/5741 = 0.45%; AD: 16/5701 = 0.28%) (Figure 2B,C). The proteins for which fewer than two peptides were detected and/or in less than 75% of the samples i.e., those considered not detected at 55 °C (NDC: 168/168+5741 = 2.8%; AD: 132/132+5701 = 2.26%) or 95 °C (NDC: 79/79+5741 = 1.4%; AD: 100/ 100+5701 = 1.72%) were low, indicating that solubilization in this one-tube reaction was optimal and independent of sample type, most likely due to the very thin tissue sections (10 µm) creating sufficient exposure to the chemical solubilization. To explore whether there are consistent protein sets that were not detected at either solubilization temperature, we selected the proteins in overlap between NDC and AD lost at 55 °C (n = 43) and at 95 °C (n = 11) for gene-set enrichment using the gene ontology tool ShinyGO [32]. No enrichment was found. Solubilization temperature does not favor specific proteins. Based on these findings, we selected 95 °C as the solubilization temperature for the subsequent synaptosome proteomics experiment (see section 2 & Supplemental protocol 1).

**Figure 2.**
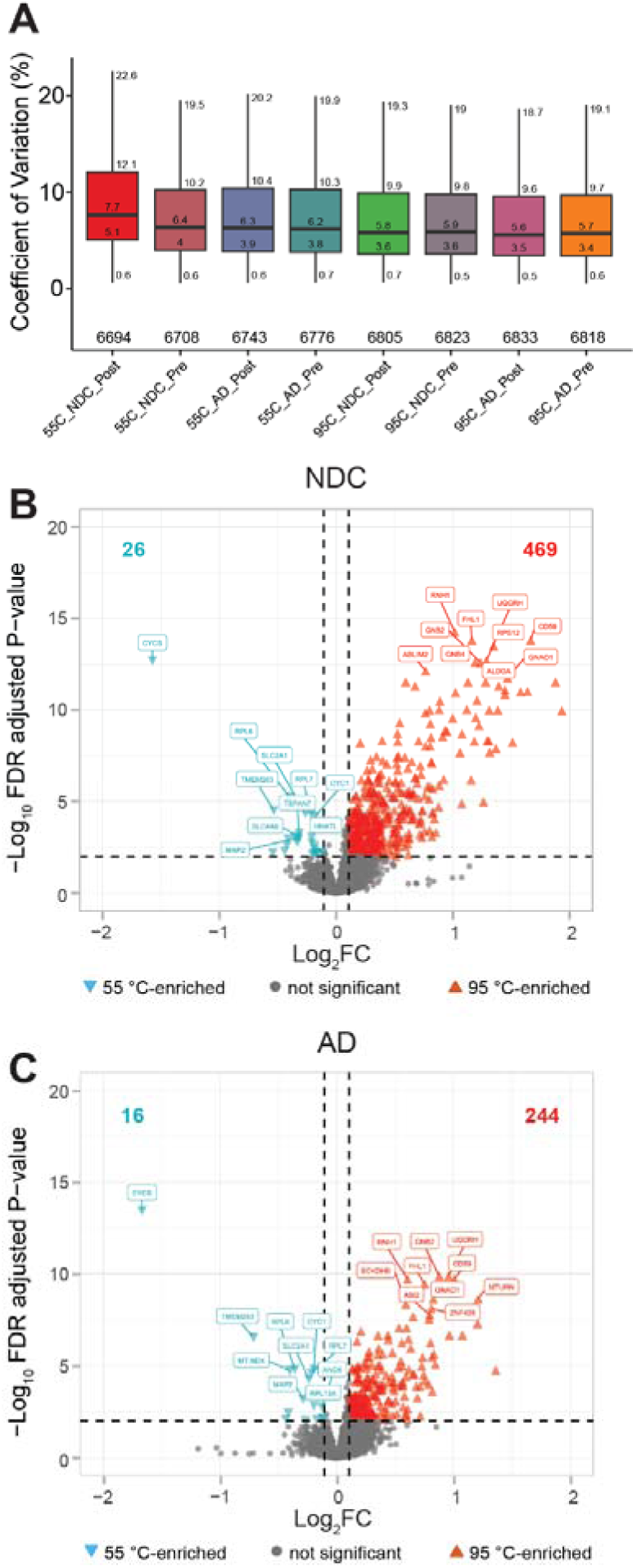
Quality control parameters for one-tube tissue lysate protocol. A) Protein-level coefficient of variation plots of all sample conditions. Values of quartiles and whiskers, and total number of proteins are indicated. A higher temperature during solubilization (95 °C) resulted in a small reduction of coefficient of variation (CoV). B,C) Volcano plot of DEqMS results showing the comparison of protein levels after solubilization at 55 °C vs. 95 °C prior to centrifugation for NDC (B) and AD (C) samples. In total, 2 tissue replicates (A, B) and 4 sample replicates (1–4) were taken per condition, resulting in 8 technical replicates per donor. Gene symbols are shown for the top 10 proteins with lowest FDR adjusted p-value (<0.01; horizontal dashed line) for both 55 °C-enriched and 95 °C-enriched. The total number of differentially expressed proteins (vertical dashed line; >|±0.10|) is indicated at the top (left, turquoise; right, red). Pre = pre-centrifugation; Post = post-centrifugation; NDC = non-demented control donor; AD = donor with Alzheimer’s Disease and comorbid schizophrenia. For a complete list of proteins, see Supplemental file 1.

Subsequent centrifugation of the samples did not lead to substantial protein loss, regardless of solubilization temperature or sample type (NDC or AD) (Figure 2). At both 55 °C (Supplemental Figure S4) and 95 °C (Figure 3), the few proteins that were less abundant due to centrifugation included collagens (95 °C: NDC: 5/5800 proteins = 0.09%; AD: 5/5816 proteins = 0.09%), while some ribosomal proteins appeared paradoxically more abundant (95 °C: NDC: 1/5800 proteins = 0.02%; AD: 3/5816 proteins = 0.05%) (Figure 3). The proteins that were not confidently detected –i.e., those considered truly “lost” proteins after centrifugation– represented a small fraction of the total protein pool (95 °C: NDC: 104/104+5800 = 1.8%; AD: 92/92+5816 = 1.6%; Supplemental file 1). Similarly, the proportion of proteins confidently detected only after centrifugation (“gained” by centrifugation) was comparable across samples (95 °C: NDC: 92/92+5800 = 1.6%; AD: 98/98+5816 = 1.7%; Supplemental file 1). Similarly, the proteins in overlap between NDC and AD not detected in the pre-centrifugation (n = 12) and post-centrifugation (n = 15) samples at 95 °C was very low, suggesting that these differences likely reflect stochastic detection of low abundant proteins rather than systematic effects of the centrifugation step. Although overall protein loss was negligible, we decided to retain the centrifugation step in the protocol. This step ensures a more representative sampling of the extracted proteins by removing potential debris resulting from the tissue collected.

**Figure 3.**
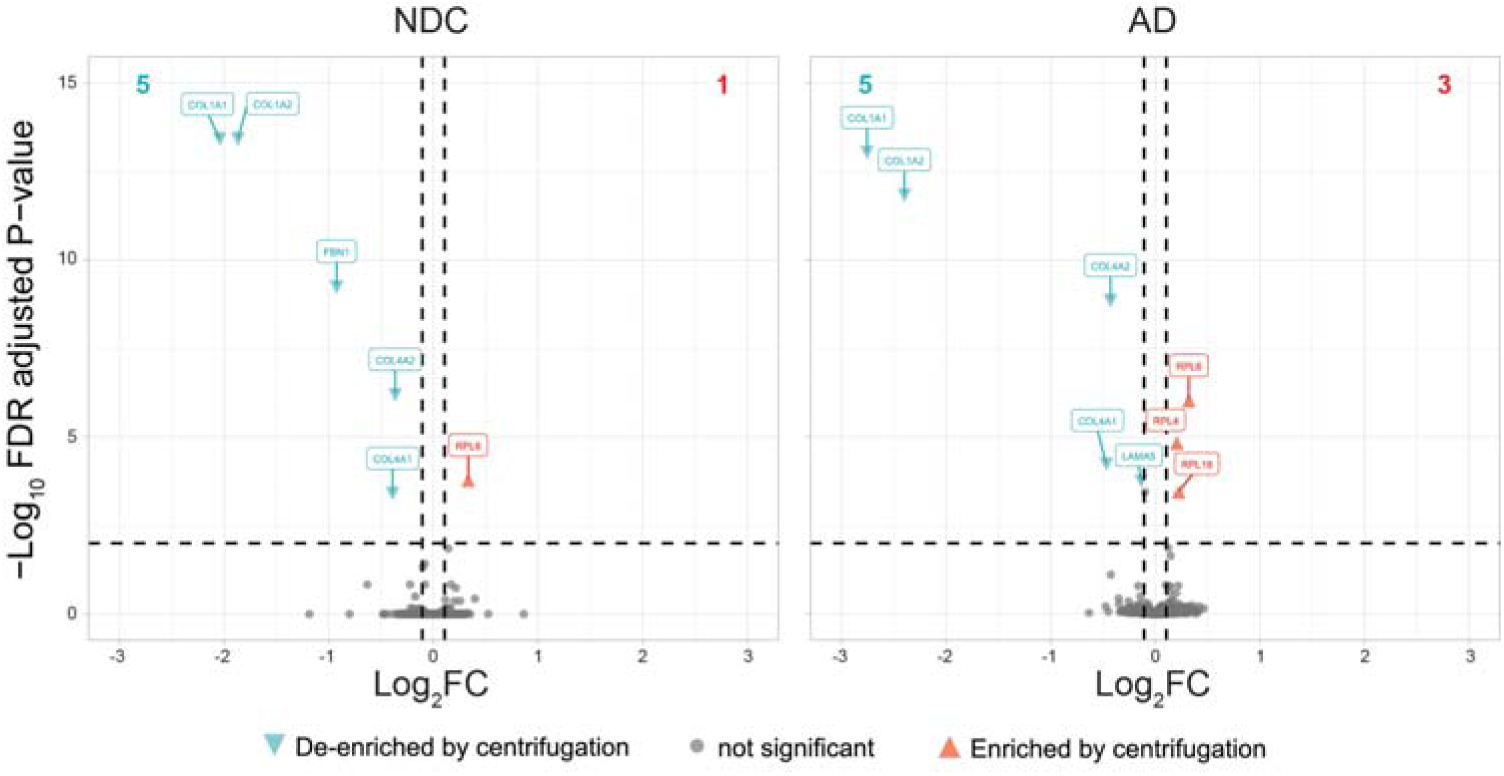
Protein (de-)enrichment by centrifugation of samples. Volcano plots showing the DEqMS contrast of pre- vs. post-centrifugation samples at 95 °C, for control samples (NDC, left) and AD samples (AD, right). Two tissue replicates (A, B) and 4 sample replicates (1–4) were taken per condition, resulting in 8 technical replicates per donor for pre- vs. post-centrifugation. Gene symbols are shown for the significant proteins that are centrifugation-depleted (turquoise) or centrifugation-enriched (red) for adjusted p-values <0.01 (horizontal dashed line). The total number of differentially expressed proteins (colored; vertical dashed line; >|±0.10|) is indicated at the top (left, turquoise; right, red). Overall, the difference in protein abundance due to centrifugation is minimal, and greatly similar between the NDC and AD samples. For a complete list of proteins, see Supplemental file 1.

The final step in optimizing our protocol involved validating the consistency of Trypsin / Lys-C digestion when processing samples of variable input level, reflecting the range of tissue weights that was observed while collecting samples from the gray matter of human tissue blocks. For this, samples from the previous experiments were pooled and subsequently diluted to simulate a series of samples without biological variability, and with a tissue weight equivalent to 10 mg, 5 mg, 2.5 mg, and 1 mg. The sTRAP workflow was performed on an equal fraction of each sample using a fixed amount of Trypsin / Lys-C (1 µg). Digestion efficiency, reported by DIA-NN as the “average missed tryptic cleavages” statistic was found to be consistent for all simulated input weights (Figure 4A), validating the use of a singular Trypsin / Lys-C condition for a wide range of protein inputs. The overall number of missed trypsin cleavage sites was low with 11.7% of all measured peptides and 9.0% of peptides detected in 100% of samples (Table S3). However, when examining the depth of the measured proteome, we observed that the number of identified peptides and proteins gradually decreased –alongside an increase in coefficient of variation– with a reduction in the sTRAP protein input (Figure 4B,C), specifically for protein inputs of 1 mg tissue (5 µg protein equivalent). This reduction in peptide and protein IDs should not originate from a reduction in total MS load, as a peptide-level normalization was performed prior to MS measurement. A possible explanation can be found in the relative increase, and therefore load contribution, of peptides that are derived from Trypsin / Lys-C itself through autolysis at higher enzyme to protein ratios. Alternatively, relative surface losses of specific peptides could become more prominent during the sTRAP workflow at lower input levels, as observed from the increase of variation in these samples. This observed variability of peptide and protein IDs across the simulated tissue levels strongly encourages the use of protein-level normalization prior to digestion (e.g. performing the BCA assay on tissue lysates) to ensure equal loading for sTRAP.

**Figure 4.**
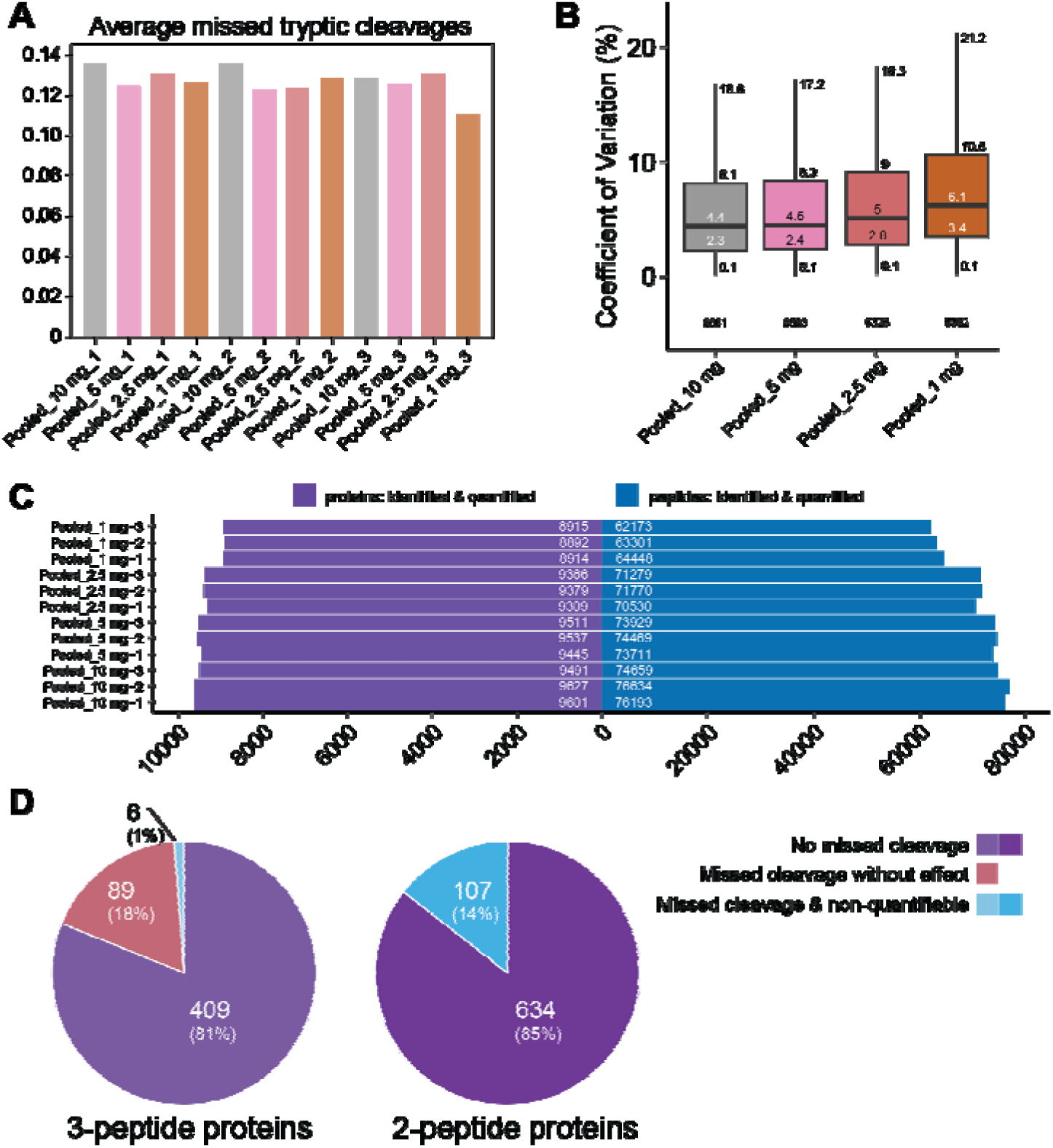
Quality controls for Trypsin / Lys-C digestion efficiency. A) Average missed tryptic cleavages of samples simulating 10, 5.0, 2.5, and 1.0 mg tissue weight input (50, 25, 12.5 and 5 µg protein equivalent for sTRAP), measured in triplicate (1-2-3), after digestion with 1 µg Trypsin / Lys-C (ratio of enzyme-to-protein = 1:5, 1:12.5, 1:25, 1:50, respectively, with 1:20 being the optimal ratio). B) Protein-level coefficient of variation of the previous samples. Values of quartiles and whiskers, and total number of proteins are indicated. C) Protein and peptide counts of all sample conditions. Number of detected peptides and proteins increased with increasing simulated input weight. D). The number of missed trypsin cleavage affecting the number of quantifiable proteins (becoming unquantifiable, blue; no effect, pink), and proteins not affected (purple) were evaluated when restricting the DIA-NN analysis from one to zero missed cleavages for proteins represented by low peptide numbers (2 and 3).

In addition, we evaluated the impact of missed cleavages on protein quantification by assessing the number of proteins that would have been unquantifiable (i.e., represented by fewer than two peptides) if zero missed cleavages were allowed (Figure 4D). This analysis specifically focused on proteins represented by a low number of peptides, as these are most susceptible to loss of quantification. We found that only a relatively small number of proteins—107 (14%) proteins represented by two peptides and 6 (1%) proteins represented by three peptides—benefited from allowing the standard DIA-NN missed-cleavage setting of one (Figure 4D).

Based on the results of our pilot experiments, we established the following optimized protocol: 1) Using samples of 2.5–10 mg tissue weight, protein solubilization was performed in 500 µL 5% SDS buffer using 2-CAA for irreversible cysteine alkylation at 95 °C for 15 minutes, with a 15-minute cool-down to 55 °C to avoid safe-lock tubes from opening due to pressure build-up. 2) Extracted proteins were centrifuged to ensure a more representative sample by removing potential aggregates. 3) Trypsin / Lys-C digestion was carried out using a fixed Trypsin / Lys-C-to-estimated protein ratio of 1:25 for samples derived from tissue weight of 5–10 mg whenever feasible. For each of the specific steps, see supplemental protocol 1. Optimal sampling is achieved by collecting multiple tissue slices to reach a total input of 2.5–10 mg, thereby ensuring balanced representation of the tissue block; however, this depends on the size and geometry of the available tissue. When only smaller amounts of tissue (e.g. 0.5–1.5 mg) can be collected, we still recommend solubilization in 500 µL, but this would result in iterative loading of the protein suspension on the sTRAP column to reach the minimal input of 12.5 µg protein (2.5 mg tissue-equivalent; *c.f.* Figure 4). The specifics of this have not been tested here.

### 3.2. Subcellular Fraction Isolation: Synaptosome Proteomics

Proteomic technologies have advanced rapidly, particularly in terms of sensitivity and quantification of proteins relative to the input material required. Subcellular fractionation –such as synaptosome isolation in neuroscience studies [37]– has been commonly employed to reduce sample complexity and enrich relevant protein subsets [38], thereby overcoming limitations in detection and quantification. However, with current mass-spectrometers now capable of identifying ∼10,000 proteins, such detection limits have become less of an issue. In this context, we evaluated whether isolating synaptosome fractions would still provide an advantage in detecting low-abundant synaptic proteins compared to whole-tissue lysates. To test this, we performed a quantitative proteomics analysis comparing whole-tissue lysates and synaptosome-enriched lysates from the same set of NDC donors within the NBB-cohort (n=8; 4 females, see Table 1). We collected 5.0 mg GM (10 µm sections) for whole-tissue analysis and 25.0 mg GM (50 µm sections). Synaptosomes were isolated using an in-house subcellular fractionation protocol (Supplemental protocol 2). The sTRAP solubilization process was adapted slightly to accommodate the distinct nature of the synaptosome samples but was otherwise followed as previously described (Supplemental protocol 1). All samples were randomized to avoid batch effects.

Although one whole-tissue lysate sample created higher CoV values for the group, excluding this sample yielded results comparable to those including the sample. Therefore, we proceeded with a DEqMS analysis using all 8 samples with age and sex as covariates, comparing whole-tissue and synaptosome groups directly. Using a relatively stringent detection limit (2 peptides per protein, 75% detection per group), we confidently measured 5816 proteins detected in both whole-tissue lysate and synaptosome preparations (Figure 5A; Supplemental file 2), of which 1379 proteins were annotated as synaptic using the synaptic gene ontology online tool SynGO (SynGO release 1.3, [33]). Note that when using DIA for quantification, peptides need to be detected across samples. For some proteins that were confidently detected in the lysate sample, they were not confidently detected after synaptosome isolation (truly “lost” proteins; Supplemental file 2). These were a small fraction of the total amount (812/(812+5816) = 12.3%), of which 27 were annotated as synaptic in SynGO (Figure 5A). Gene-set enrichment using the gene ontology tool ShinyGO [32] for these 812 proteins showed they belonged primarily to the nucleus (Supplemental Figure S6), which was expected based on the way synaptosomes were isolated (See Supplemental Protocol 2). Proteins not confidently detected in the lysate, but well detected in the synaptosome fraction, were only 248 (248/(248+5816) = 4.1%), of which 33 were annotated as synaptic in SynGO. Gene-set enrichment of these 248 proteins did not show overrepresentation for any cellular component term, suggesting these proteins were from diverse origin. Because we used stringent criteria for quantification (i.e., 2 peptides and presence in 75% of the samples), we checked what the effect would be of lowering the detection limit to 1 peptide. From the 248 undetected proteins in the lysate, 180 proteins could then be identified, while 68 proteins were not detected at all in the lysate samples. To check whether this was just a matter of stochasticity, we plotted the frequency of peptides detected in the synaptosome fraction (Supplemental Figure S5). From these 68 proteins, 50 were detected with 2 peptides in the synaptosomes, and 18 with ≥3 peptides, indicating that this might well have been a matter of chance detection in the synaptosome fraction, similar to the many proteins detected only by 1 peptide in the lysate. Yet, a small set of proteins that was absent or expressed at very low levels in the lysate (0 to 1 peptide) seems highly enriched in the synaptosome fraction (detected with ≥4 peptides in synaptosomes; Supplemental Figure S5). Albeit that gene-set enrichment did not show overrepresentation in cellular component, these proteins represent a diverse group of molecules with (possible) roles in synaptic organization, signaling, and membrane dynamics. Several (e.g., GRIP2, GABRG3, SLC17A8) are established synaptic or neurotransmission-related proteins [39–42], or the AMPA receptor auxiliary subunit SHISA6, which is highly expressed in the dendrites of the hippocampus CA1 region [43]. All are present in the SynGO database, reinforcing the specificity of our synaptosome preparation. Others (PRKCQ, TYK2, LRRK1, PTPN13 and XPR1) are kinases, phosphatases and phosphate regulators that may regulate synaptic plasticity or local signaling cascades [44–48]. Proteins such as LRRK1, RAB27A, and SPIRE2 [49–53] suggest active vesicular trafficking, and cytoskeletal remodeling at the synapse. The presence of UNC5D, PDZRN3, PTPN13 and N4BP3 indicates potential involvement in synaptic function, or facilitating cell communication, as all have been implicated in neuronal migration [54–57].

**Figure 5.**
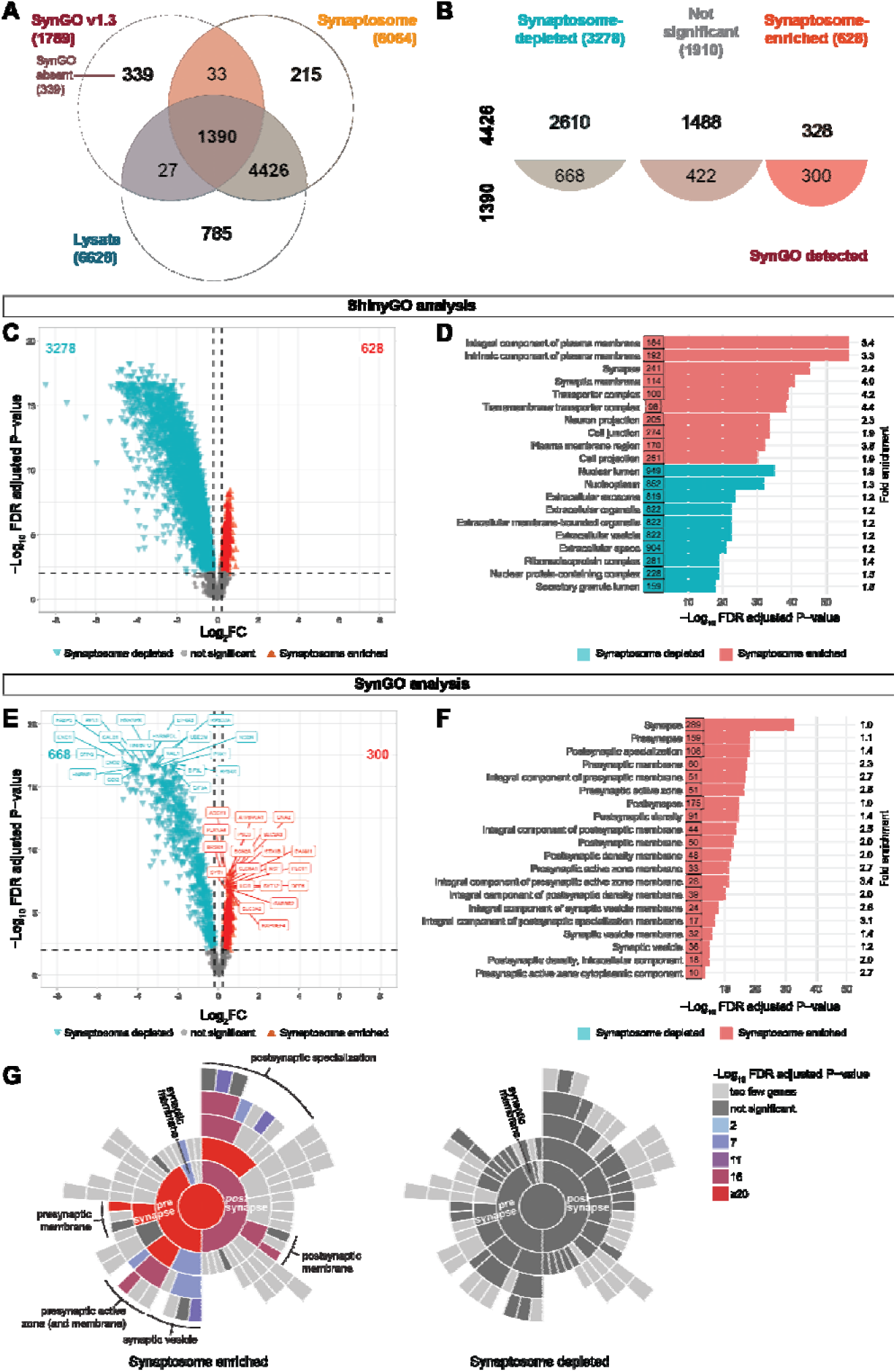
Enrichment of synaptic proteins in the synaptosome fraction. A) Venn diagram showing the total number of detected proteins and the proportion of SynGO annotated synaptic proteins (“SynGO detected”) and SynGO proteins not detected in whole-tissue lysate (“Lysate”) and synaptosome (“Synaptosome”) samples. B) Diagram showing the division of proteins of the contrast of whole-tissue lysate vs. synaptosome samples (Synaptosome-depleted, turquoise; Synaptosome-enriched, red; Not significant, gray; see panel C), and whether or not the proteins were annotated in SynGO (1390 and 4426 resp. in panel A. C) Volcano plot showing the contrast between whole-tissue and synaptosome for NDC donors (n = 8 for whole-tissue vs. n = 8 for synaptosomes). The largest group of proteins shows depletion (turquoise) in the synaptosome samples, while a smaller number of proteins is enriched (red). The total number of differentially expressed proteins is indicated at the top (left, turquoise; right, red). D) Bar graph showing the top 10 overrepresented GO annotations for cellular component of synaptosome enriched and depleted proteins, using ShinyGO. The number of proteins involved in each pathway as well as the fold enrichment are indicated. E) Volcano plot of the same contrast as C) but showing only the proteins annotated in SynGO. The top 20 proteins on each side are labeled. F,G) SynGO GSEA of synaptosome enriched and depleted proteins (see E) in terms of bar graph showing the top 20 overrepresented SynGO annotations of synaptosome enriched proteins (F), or sunburst plots (G). Synaptosome depleted proteins did not show significant overrepresentation in SynGO; (see Supplemental Figure S5 for ShinyGO annotation). SynGO sunburst plots showing the overrepresentation of synaptosome enriched and depleted proteins for cellular component in SynGO (G). For a complete list of proteins, see Supplemental file 2.

Next, we checked to what extent curated synaptic (i.e., SynGO) proteins were not detected in either lysate or synaptomes. From the entire SynGO database of 1789 curated protein entries [33], 339 proteins were not detected in either sample (‘SynGO absent’ proteins (*cf*. Figure 5A). Gene-set enrichment of this group of proteins indicated that among the highly overrepresented GO groups were acetylcholine receptor subunits, as well as receptor proteins from several families (Supplemental Figure S6, Supplemental file 2). The expression level of these proteins is very low, and levels of acetylcholine receptor subunits typically have been measured by receptor binding studies with subsequent immunoprecipitation [58,59].

From a quantitative perspective, our comparative LC-MS/MS analysis between whole-tissue lysate vs. synaptosome showed the significant (adjusted P-value <0.01) enrichment of a relative small set of proteins in synaptosomes (628 out of 5816 = 10.8%), and a marked depletion of a relative large set of proteins (3278 out of 5816 = 56.4%) (Figure 5B,C). As expected, synaptic terms were enriched in proteins with a higher expression in synaptosomes, and nuclear and extracellular proteins were enriched for proteins with a higher expression in the whole-cell lysate samples (Figure 5C). From synapse-enriched and synapse-depleted proteins, 47.8% (300 out of 628) and 20.4% (668 out of 3278) were annotated as synaptic proteins, respectively (Figure 5E). Specifically focusing on the detected SynGO proteins in this contrast (Figure 5E; 300 enriched & 668 depleted), we observed enrichment of typical synaptic proteins, such as proteins of the vacuolar ATPase complex (e.g. ATP6V0A1) [60] and metabotropic GABA_B_ receptor subunits (e.g. GABBR2) [61], and synaptic depletion of heterogeneous nuclear ribonucleoproteins (e.g. HNRNPK and HNRNPL) [62]. Further gene-set enrichment using ShinyGO [32] showed that proteins enriched in synaptosomes were predominantly associated with membrane components (ion channels, transporters) and synaptic compartments (Figure 5D). In contrast, proteins enriched in whole-tissue lysates (i.e., synaptosome-depleted) were overrepresented in categories such as cytosolic ribosomes, nuclear related components, and extracellular components (Figure 5D). A more in-depth examination of synapse-specific proteins using SynGO showed a significant overrepresentation in both presynaptic and postsynaptic compartments for the 300 synaptosome enriched SynGO proteins (Figure 5F,G). Although many SynGO proteins were detected in the synaptosome depleted set, these 668 proteins did not show significant enrichment for cellular localization terms in SynGO (Figure 5G). GO annotations of this synapse-depleted SynGO protein set indicated predominant associations with ribosomal components and organelles of extracellular origin (Supplemental Figure S6), as required for local translation to meet the plasticity demand of the synapse [63,64]. Together, these proteins are not classical synaptic proteins (e.g., glutamate receptors), rather are (multifunctional) proteins with multiple cellular localizations, one of which is the synapse.

Comparing whole-lysate and synaptosome samples is inherently challenging, as the different nature of these preparations may complicate normalization and raise concerns about potential bias, as reflected by the skewness of the volcano plot. To address this, we analyzed the distribution of SynGO-annotated proteins across 50-protein bins ordered by log2 fold change (Supplemental Fig. S7). Given the high number of SynGO proteins in bins 1–6, the remainder of the proteome was divided into three additional bins of comparable size, plus a final residual bin. Gene set enrichment analysis (GSEA) of SynGO terms revealed a clear overrepresentation in bins 1–6 and, to a lesser extent, in bins 7–18, the latter approaching the threshold for differential expression. Based on these results, we conclude that the observed proteomic shifts are driven by genuine compartment-specific protein enrichment. Furthermore, the normalization approach implemented in MS-DAP [30] is sufficiently robust to accommodate large differences between the sample types analyzed.

Previous correlation profiling of subcellular fractions from mouse hippocampus, cortex, and cerebellum—including microsomes, synaptosomes, synaptic membranes, and postsynaptic densities—revealed consistent patterns of functional grouping and subcellular localization [19]. Based on these findings, we selected sets of mouse proteins that were previously identified as either enriched (n = 192) or depleted (n = 134) in synaptosomes and synaptic membranes from the cortex. Of these, we detected 79 synapse-enriched and 117 synapse-depleted orthologs in our human cortex proteomics dataset. Notably, many of these proteins exhibited similar relative abundance patterns in the human samples, with a strong and statistically significant correlation between mouse and human data (Pearson r = 0.64, P < 0.001; Figure 6A). Among the human proteins that mirrored the synaptic enrichment or depletion observed in mice (79 and 117 proteins, respectively), functional annotation revealed localization to either pre- and post-synaptic compartments (synapse-enriched) or to cytosolic, ribosomal, or membrane-associated structures (synapse-depleted) (Figure 6B,C), similar to the previous analysis shown in Figure 5 and Supplemental Figure S6. Three notable subsets among the synapse-depleted proteins were the relatively high enrichment of the postsynaptic (intermediate filament) cytoskeleton and neurofilament groups, which included proteins like neurofilament medium chain (NEFM), and neurofilament light chain (NEFL) (see Supplemental file 2). They are known to function in the postsynaptic compartment [65,66], and are indeed enriched in the postsynaptic density fraction, but not in the synapse fractions [19]. We found these cytoskeletal proteins consistently identified with high confidence, each supported by more than 49 unique peptides, and they were equally detected across both sample types. Hence, the observed synaptosome-depletion stems from differences in relative abundance and subcellular localization (post-synaptic density-enriched) rather than technical variability. Surprisingly, a part of the proteins that were enriched in the synaptosome fraction in mice showed opposite regulation in our human dataset (n = 54). Gene ontology analysis showed that these proteins belonged to vesicle pathways and the Golgi apparatus (Supplemental Figure S8).

**Figure 6.**
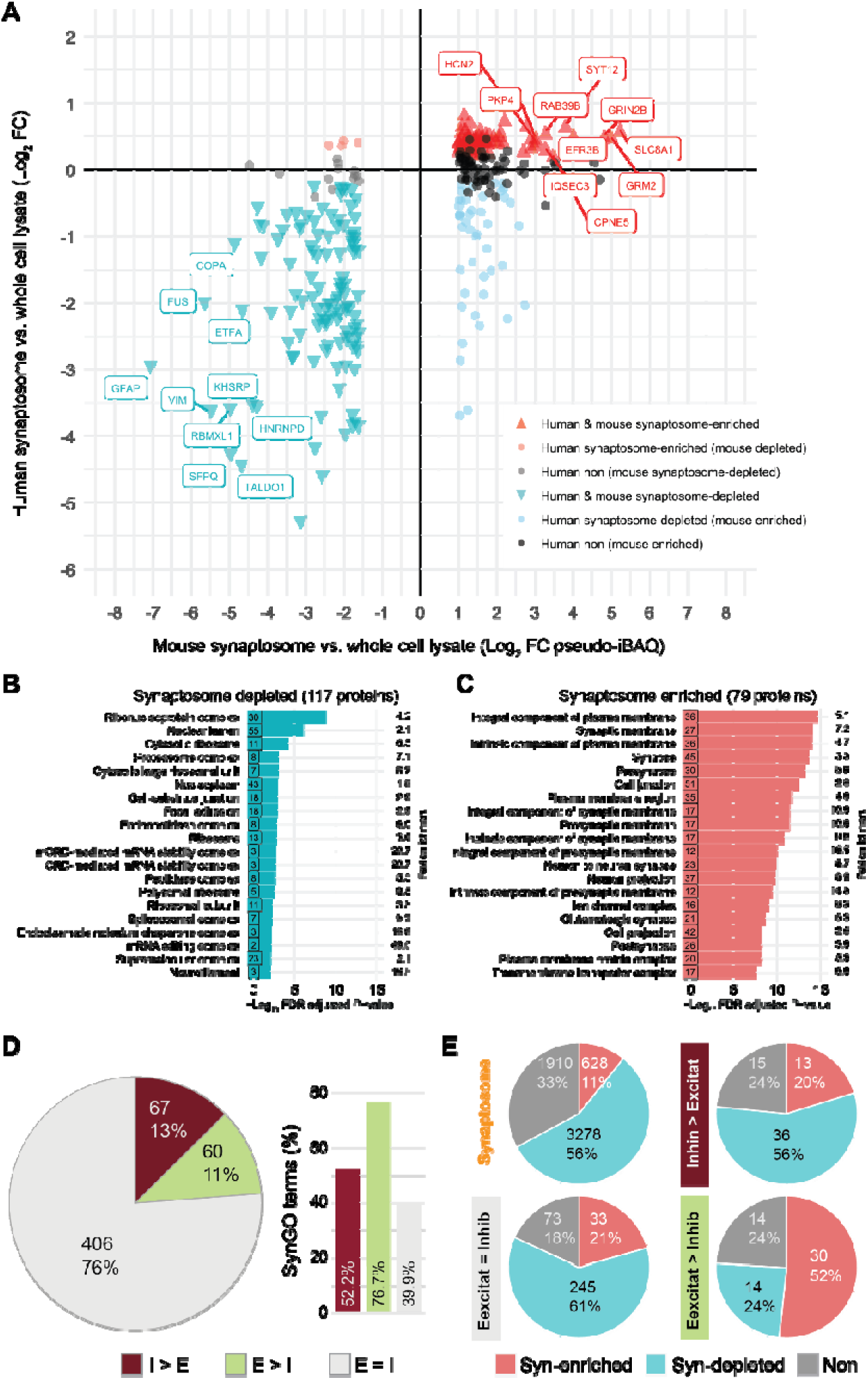
Cross-validation of synaptosome enrichment and depletion. A) Using a previously published dataset of mouse cortical synapse-enriched and depleted proteins by differential biochemical fractionation [19], selected proteins (n = 326) were retrieved from our human NDC lysate vs. synaptosome data set, and their relative ratios (log2) were compared. The top 10 proteins with the highest enrichment in both mouse and human synapses are labeled. B,C) Bar graph showing the top 20 overrepresented GO annotations of mouse synaptosome (pseudo-iBAQ values) and human synaptosome proteins that were both depleted in synaptosomes (B) and enriched in synaptosomes (C) using ShinyGO. The number of proteins participating in each annotation, as well as the fold enrichment are indicated. D) Pie chart showing distribution of proteins related to inhibitory (I>E), excitatory (E>I) cells and those with no preference (E=I) based on the van Oostrum et al. dataset [68], and bar plot showing their % SynGO v1.3 annotations. E) Pie charts showing the distribution of synaptosome-enriched, synaptosome-depleted and non-regulated protein terms as detected in the synaptosomes in comparison to that of the markers for the 3 cell types. Numbers and % of total are indicated. For a complete list of proteins in each analysis, see Supplemental file 2.

Next, we compared our data with those of a meta-analysis of the synaptic proteome, covering 8044 unique protein IDs detected from 58 published synaptic proteomic datasets derived from different parts of the human, mouse and rat brain by Sorokina et al. [67] (Supplemental Figure S8). Overlap with our synaptosome dataset was slightly higher (5096 proteins, 84.0%) than that for the lysate dataset (5414 proteins, 81.7%). SynGO proteins were represented at a similar rate in the Sorokina dataset (1522, 18.9%), compared to that of the lysate (1298, 19.6%), and were both at a slightly lower rate than the synaptosome (1302, 21.5%) samples. Remarkably, the number of proteins that were unique to the Sorokina dataset was significantly higher (2363, 29.4%) than that for the synaptosome (109, 1.8%) and lysate (357, 5.4%) set. This could reflect biological variation due to use of data from different species and brain regions. However, a ShinyGo overrepresentation analysis of the 2363 unique proteins indicated a large proportion of proteins related to the nucleus (chromosome/chromatin, nucleosome etc.) (Supplemental Figure S8). This suggests that, at least in part, contamination due to the type of (biochemical) isolation used contributed to annotating these proteins as synaptic.

Lastly, we compared our data to a proteomics dataset where the authors did not only isolate synaptosomes but also used fluorescent labeling in transgenic mice to find unique synapse-type-enriched proteins [68]. To ensure we selected synaptic proteins from the cortex, we took those that showed a significant enrichment for synapsin1 after sorting [69]; this resulted in 533 proteins (Supplemental file 2). Subsequently, we divided these proteins based on the enrichment of the Camk2 vs. GAD labeling, as being excitatory-enriched (E>I; 60, 11.3%), inhibitory-enriched (I>E; 67, 12.6%), or common (E=I; 406, 76.2%) (Figure 6D). As the authors mentioned [69], these proteins show a high percentage of SynGO annotations, and this is specifically true for the excitatory-enriched proteins (E>I; 46, 76.7%; Figure 6D). From the 533 synaptic proteins, we found a high coverage in our samples, with 523 detected in the lysate and 524 detected in the synaptosome samples, and 523 proteins in overlap (quantified). Inhibitory-enriched as well as common marker proteins show a distribution of synaptosome-enriched, -depleted, and non-regulated proteins similar to that observed in the total population (Figure 6E; *c.f.* Figure 5C), albeit with a slightly higher proportion of synaptosome-enriched proteins at the expense of non-regulated proteins. Remarkably, the excitatory-enriched marker proteins show a significant increase (Chi-2 (12, n = 581) = 25.6, P<0.001) in synaptosome-enriched proteins vs total proteome markers, mostly at the expense of synaptosome-depleted proteins (Figure 6E). This indicates that our synaptosome-enriched proteins largely stem from excitatory neurons, which are abundantly present in the cortex.

## 4. Discussion & Conclusion

In this study, we first optimized our pre–mass spectrometry (pre-MS) workflow for neuroproteomic analysis with LC-MS/MS of post-mortem human grey matter (GM) and then used this optimized pipeline to compare whole-tissue lysate and synaptosome-enriched proteomes from human cortex. Post-mortem brain samples are typically received as frozen tissue blocks and are often processed for multiple downstream applications, including proteomics and immunohistochemistry. In this context, the separation of GM from white matter (WM) provides a critical advantage. GM is enriched in neuronal cell bodies and synaptic components [5], whereas WM contains abundant myelin proteins and oligodendrocyte-derived material [6] [7]. Analyzing GM separately improves the detection of lower-abundant neuronal and synaptic proteins that could otherwise be masked by highly abundant myelin proteins, thereby increasing the biological specificity and interpretability of proteomic data.

Laser capture microdissection from thin tissue sections (5–10 µm) provides a precise approach to GM isolation [70,71], but its labor-intensive nature makes it impractical for large-scale proteomic studies (e.g. >200 samples). To balance specificity and scalability, we optimized a workflow based on free-hand dissection of GM from thin cryosections to generate whole-tissue lysates. Importantly, the use of thin (10 µm) sections substantially increases the accessible surface area of the tissue, facilitating efficient chemical solubilization. We further demonstrated that elevated solubilization temperatures markedly enhanced protein extraction efficiency, underscoring the importance of optimizing tissue processing parameters for human brain samples.

### Optimization of tissue solubilization and digestion

Tissue solubilization is a critical preparatory step in any proteomic workflow, as it directly influences protein yield, integrity, and the breadth of proteome coverage. Two principal strategies are commonly employed: mechanical and chemical solubilization [18]. Mechanical disruption methods, such as tissue grinding, bead beating, and sonication [8,72–74], are widely used to break down the structural integrity of tissues and cellular membranes and are widely used in subcellular proteomics, such as organelle or synaptic compartment isolation [75,76]. However, these approaches can introduce batch-to-batch variability and are sensitive to technical inconsistencies, particularly when working with small or heterogeneous human samples [2,72,77].

In contrast, chemical solubilization relies on the use of surfactants (e.g., SDS, Triton X-100, CHAPS) and chaotropic agents (e.g., urea, thiourea, guanidine hydrochloride) to denature and solubilize proteins, including membrane-associated and hydrophobic proteins that are typically underrepresented in proteomic datasets [78]. The use of well-defined buffer compositions improves reproducibility and facilitates scalability, making chemical lysis particularly well suited for high-throughput proteomic pipelines [79]. A key limitation of this approach is the need to remove MS-incompatible reagents prior to LC-MS/MS analysis, which can be effectively achieved through protein precipitation or filter-based methods such as sTRAP [3].

We here further optimized the sTRAP digestion protocol previously published by our lab [3], employing an alternative chromatographic column that is substantially more cost-effective. The rational for this was to accommodate the specific requirements of human GM samples, including tissue quantity and protein input. First, we employed a high extraction temperature (95 °C) in a one-tube workflow, which benefits from a large surface area–to–volume ratio. While the original Protifi S-TRAP protocol recommends vortexing at room temperature or the inclusion of sonication or benzonase steps followed by reduction at 55 °C, our data demonstrate that elevated extraction temperature markedly enhances protein solubilization and identification. This effect was likely further strengthened by using thin gray-matter tissue sections, which facilitate more efficient heat transfer and extraction of insoluble proteins. Second, we substituted the vendor-recommended S-methyl methanethiosulfonate (MMTS) with 2-chloroacetamide (2-CAA). The use of 2-CAA enables complete and irreversible cysteine alkylation, rendering modified cysteines inert to TCEP and thereby improving reproducibility, peptide identification (*c.f.* Table S2), and confidence in peptide-spectrum matches, while also simplifying database searching [35]. Importantly, 2-CAA is compatible with a wide range of buffers, including SDS, which we used here to achieve more efficient solubilization of brain tissue and membrane-associated proteins. These adaptations ensured a highly standardized and efficient sample preparation workflow, which is an essential prerequisite for detecting subtle biological effects in human post-mortem tissue. Importantly, these optimizations did not lead to selective loss of specific protein groups, as evidenced by the low overlap among non-detected proteins across conditions. Together, these modifications facilitate the analysis of larger sample cohorts, thereby enhancing the scalability and applicability of the workflow.

Besides sTRAP –offering a practical combination of affordability, straightforward design, and high reproducibility [80]–, numerous other sample preparation techniques have become available for proteomics (see Box 1), some of which were not published at the time of the experiments performed in the current study.

An important consideration throughout the entire experimental process that should be practiced independent of using rodent or human tissue as input material – including tissue dissection and LC-MS/MS analysis– is a strict standardization and randomization [81]. This is especially important in large-scale studies where experiments are spread out over extended periods. To mitigate confounding batch effects in the interpretation of the resulting data, we evenly distributed the samples of each experimental group across all experimental batches for every experimental step. In addition, we randomized the order of sample dissection, sTRAP preparation, and LC-MS/MS analysis to avoid confounding effects. Notably, we observed a trend in LC-MS/MS analysis where the order of samples ran affected the number of identified peptides (*c.f*. Supplemental Figure S1). This underscores the critical role of rigorous standardization and randomization in large-scale proteomic workflows to ensure that technical variation affects all experimental groups equally. The workflow described here may serve as a practical guide for researchers aiming to standardize and refine neuroproteomic analyses from human brain tissue dissection through LC-MS/MS analysis.

In the present study using human brain samples, we confidently identified ∼6,000 to over 7,400 proteins from human brain tissue, based on an initial detection of more than 82,000 to ∼105,000 peptides (Figures 2 and 5). This depth exceeds those reported in comparable studies analyzing whole-tissue lysates from human cortex or brain organoids [82–84], and approaches that of previous studies using pre-fractionation of isolated synaptosome samples [Table S1 [16]). Together, these results reflect recent advances in mass spectrometry sensitivity and support the suitability of chemically solubilized whole-tissue lysates for high-depth human neuroproteomics.

##### Box 1: Alternative methods for proteomics sample preparation

***NanoPOTS (nanodroplet processing in one pot for trace samples)*** *performs all the proteomic processing steps (extraction, alkylation, digestion and peptide collection) in a semi-automatic manner within a single droplet, making this technique ideal for small biological samples (as low as 10-100 cells) as it reduces sample loss and increases robustness and reproducibility for high-throughput studies [39]. However, it requires highly specialized equipment and costly materials, making this technique less accessible*.

***SP3 (single-pot, solid-phase-enhanced sample-preparation)*** *uses hydrophylic magnetic beads to capture proteins in a single tube, facilitating addition and removal of reagents needed for protein preparation for mass spectrometry and resulting in virtually no loss of material [40]. SP3 is compatible with a wide range of reagents and can be used for both low and high quantities of input material, but with protein amounts of >500* μ*g –as in the current study– the beads show aggregation, sticking to pipette tips and tube walls and resulting in protein loss*.

***iST (in-StageTip)*** *is performed in a single device as well, a pipette tip with reaction chamber in which all preparation steps take place, and a C18 filter at the bottom for final peptide cleanup [41,42]. It is a simple and robust method, but it shows incompatibility with detergents such as SDS that greatly improve cell lysis and protein yield [40]*.

*The **One-Tip** methodology is a more recent development, building upon the one-pot reaction [43]. Requiring only two pipetting steps, it combines a cell lysis and digestion buffer with the sample suspended in PBS into an Evotip (Evosep Biosystems) for pre-MS processing. It is suitable for single cell analysis, but also for bulk proteomics from tissue sections*.

### Whole-tissue versus synaptosome proteomics

In the second part of this study, we directly compared human whole-tissue and synaptosome-enriched samples to assess their relative strengths in detecting disease-relevant protein changes. We found that MS sensitivity was sufficient to detect most of the low-abundant (synaptic) proteins in whole-tissue lysates (*c.f.* Figure 5, Supplemental Figure S5). We observed 1417 SynGO proteins in whole-tissue lysate (21.4% of proteome), and 1423 in isolated synaptosomes (23.5% of proteome). Moreover, virtually all SynGO proteins detected in our synaptosome fraction were also present in the whole-tissue comparison (97,8%). Given the advancement of MS sensitivity, isolation of synaptosomes is not required anymore when considering <u>detection</u> of synaptic proteins. In addition, synaptosome preparation is inherently more time-consuming, labor-intensive, and hence leads to a higher between-sample variability, which may limit its practicality in large-scale studies. However, subcellular fractionation retains clear biological value because it allows the detection of compartment-specific regulation that is likely masked in a whole-tissue context. Thus, combining lysate-level and synaptosome-level proteomics provides a more complete and biologically informative picture of synapse-specific processes and their modulation, revealing compartment-resolved regulatory mechanisms.

Based on these considerations, we propose whole-tissue proteomics as an effective first-line strategy to identify disease-associated proteomic changes. This approach offers a broad, rapid, and technically robust overview of molecular changes without compromising the detection of (low-abundant) synaptic proteins. Whole-tissue analysis is particularly advantageous when tissue is collected in parallel for both whole-tissue and synaptosome preparations as described here. In such cases, initial whole-tissue data can highlight key biological processes or pathways that may warrant deeper, targeted (synapse-focused) investigation.

### Synaptosome isolation and interpretation

Synaptosomes in this study were isolated using a sucrose density gradient [26], a method with which our laboratory has extensive experience for synaptosome, synaptic membrane, and postsynapse enrichment [19,24,76,85,86]. Alternative gradient-based approaches, including Ficoll (sucrose and epichlorohydrin copolymer, [26,87] and Percoll (colloidal silica particles covered in polyvinylpyrrolidone monolayer, [88]), are also commonly used. While these methods share similar principles, sucrose density gradients have been shown to result in lower contamination by extrasynaptic mitochondria than Ficoll gradient [75]. Although Percoll gradients offer increased speed [88]), sucrose gradients yield a higher proportion of viable synaptosomes and preserve synaptic vesicles more effectively [89], while also reducing mitochondrial contamination compared to Ficoll gradients [75]. Together, these considerations support the choice of sucrose density gradient fractionation for the present study.

It is important to emphasize that biochemical fractionation enriches rather than purifies subcellular compartments and therefore carries an inherent risk of co-isolating non-synaptic proteins. A limitation of our study is the lack of independent assessment of synaptosome purity or structural integrity (e.g., by electron microscopy), although this method has previously been validated in our laboratory ([24], Dr. Rao-Ruiz, personal communication). Residual contamination from non-synaptic membranes and organelles is therefore unavoidable and should be interpreted accordingly [9,90], as also reflected by the presence of nuclear contaminants in the synaptic proteome meta-analysis (*c.f.* Figure 6E [67]).

Despite these limitations, multiple lines of evidence support the biological relevance of our synaptosome-enriched dataset. Annotated synaptic proteins (SynGO; [33]) were significantly enriched in the synaptosome fraction and showed strong overrepresentation in gene ontology pathways associated with both presynaptic and postsynaptic compartments (*c.f.* Figure 5). Proteins relatively depleted from the synaptosome fraction did not exhibit enrichment for synapse-specific pathways, and proteins absent from synaptosomes were predominantly nuclear—consistent with their removal during early fractionation steps. Using our optimized workflow, we could confidently identify and quantify over 6,600 proteins in whole tissue lysates, exceeding coverage reported in comparable human studies [8,71,91].

In synaptosome-enriched samples, we identified and quantified over 6,000 proteins. Although this depth is comparable to the 2019 study of Alzheimer donors by Hesse et al. (Table S1; [17]), only 440 (8.2% of proteome) of these were annotated as SynGO proteins. Several factors may contribute to this relatively low proportion. First, methodological differences in synaptic enrichment—such as the use of crude P2 fractions instead of gradient-purified synaptosomes—can substantially affect synaptic specificity. Second, disease-related synapse loss or protein depletion may selectively reduce the detectability of synaptic proteins in neurodegenerative cohorts. Indeed, in the study by Plum et al. [15], which examined synaptosome proteomes from Parkinson’s disease and control donors (Table S1), SynGO coverage reached 21% overall but this increased to 42.2% when considering control donors alone. In contrast, Kandigian et al. [92] reported similar SynGO coverage in control (35.4%) and Alzheimer’s disease (33.7%) samples (Table S1), using differential (ultra)centrifugation without gradient purification.

Importantly, higher proportions of SynGO proteins are often observed in studies with more limited overall proteome coverage (Table S1), where the most abundant and well-annotated synaptic proteins are preferentially detected, see for example the study of Kumar et al. [93] . Conversely, studies achieving very deep proteome coverage tend to report lower relative SynGO percentages. This is exemplified by the large-scale synaptic proteomics study by Aryal et al. (>7,200 quantified proteins; Table S1 [16]), in which SynGO proteins accounted for only 19.5% of the dataset; very similar what we reached here (23.5%). Together, these comparisons suggest that the proportion of SynGO-annotated proteins is strongly influenced by both the depth of proteome coverage and the specific enrichment strategy used, rather than serving as a simple indicator of synaptic specificity.

### Proteome coverage and future directions

Despite the depth of proteome coverage reached, our data also highlight current limitations in human brain proteomics. Transcriptomic analyses indicate that more than 14,000 protein-coding genes are expressed in the human cortex [94] [95], implying that a substantial fraction of the proteome remains undetected. This is likely due to the complexity of human brain tissue and the current sensitivity limits of LC-MS/MS. Strategies to reduce sample complexity—such as optimized acquisition modes, physicochemical fractionation, or subcellular enrichment [92] [96,97]—as well as deeper proteomic approaches using multiple proteases and fragmentation methods [16,98,99], may further expand coverage in future studies.

On the other hand, increased MS sensitivity also raises the likelihood of detecting non-synaptic proteins in synaptosome preparations [67] (*c.f.* Supplemental Figure S7), complicating interpretation in human tissue where genetic labeling strategies are not available [68,100].

### Conclusions and recommended strategy

Although coverage of synaptic proteins in our whole-tissue proteomics dataset was largely comparable between that of synaptosome-enriched samples (see Figure 5), subtle or compartment-specific synaptic protein regulation may be obscured in whole-tissue analyses, particularly for proteins expressed in multiple cellular compartments. Conversely, synaptosome proteomics introduces challenges related to normalization and interpretation when synapse number itself is altered, as in neurodegenerative conditions [11,101].

Therefore, we recommend a tiered strategy: initiate analysis with whole-tissue proteomics to identify disease-associated processes and follow up with synaptosome proteomics on pre-dissected samples for both analyses to gain deeper insight into synaptic composition and population-level changes, while maintaining the same set of donor samples. This combined approach maximizes sensitivity, specificity, and interpretability across both cellular and subcellular levels and provides a robust framework for future large-scale human neuroproteomics studies. The gain of this two-tiered strategy is however limited and may –depending on the disease type– not be worth the effort.

## Supporting information

Supplemental Protocol 2_Synaptosome isolation

Supplemental Protocol 1_Pre-MS workflow

## Supplementary Materials & data availibility

The following supporting information is available:

Supplemental file 1 contains quantification results for the pilot proteomics experiment (relates to Figures 1–3), testing temperature, centrifugation and trypsin concentration on NDC and AD samples

Supplemental file 2 contains quantification results for the whole-tissue lysate vs synaptosome proteomics experiment (relates to Figures 4 & 5), on NDC samples;

Supplemental protocol1, contains the step-by-step optimized pre-MS protocol for whole-tissue lysate samples (tissue sections cut at 10 µm);

Supplemental protocol2, contains the step-by-step optimized protocol for synaptosome isolation from 50 µm tissue sections .

## Author Contributions

Conceptualization, FRK, RVK, and SS; Methodology, FRK, RVK, TSZK; Validation, FRK, RVK, ABS and FTWK; Formal Analysis, FRK, RVK and FTWK; Investigation, FRK; Data Curation, SS; Visualization, FRK, ABS and SS; Writing – Original Draft Preparation, FRK, TSZK and SS; Writing – Review & Editing, FRK, FTWK, RVK, TSZK, ABS and SS; Supervision, ABS and SS; Funding Acquisition, FRK and SS. All authors have read and agreed to the published version of the manuscript.

## Informed Consent & Ethical approval

Informed consent was obtained from all donors and their next-of-kin for the use of material and clinical data for research purposes, as outlined in https://www.brainbank.nl/about-us/ethics/. The NBB autopsy procedures and the use of tissue for research were approved by the Ethics Committee of Amsterdam UMC (registered with the US Office of Human Resource Protections as IRB00002991 under Federal wide Assurance number 00003703), location VUmc, Amsterdam, The Netherlands, albeit that according to the Dutch Act on Medical Research Involving Human Subjects (WMO) no ethical approval is required for collection nor use of donated brain tissue.

## Acknowledgments

This work would not have been possible without the help of Berna Özer, who assisted in sample preparation across experiments. Brain tissue was made available by Inge Huitinga, head of the Neuroimmunology group at the Netherlands Institute for Neuroscience, Amsterdam. We would also like to thank Dennis Wever (Huitinga group); his expertise in the free-hand dissection of GM was essential for the tissue collection. During the preparation of this manuscript, the authors used ChatGPT (OpenAI, GPT-4&5) for text editing. The authors have reviewed and edited the output, and they take full responsibility for the content of this publication.

## Abbreviations

The following abbreviations are used in this manuscript:

LC-MS/MS: Liquid Chromatography-tandem Mass Spectrometry
sTRAP: suspension TRAPping
NBB: Netherlands Brain Bank
SFG: Superior Frontal Gyrus
NDC: Non-Demented Control
AD: Alzheimer’s Disease
GM: Gray Matter
PMI: Post-Mortem Interval
Braak LB: Braak Lewy Bodies
APOE: Apolipoprotein E
SDS: Sodium Dodecyl Sulfate
BCA: Bicinhoninic Acid
DEqMS: Robust statistical inference for quantitative LC-MS proteomics
DIA-NN: Data-Independent Acquisition - Neural Networks
dia-PASEF: Data-Independent Acquisition - Parallel Accumulation Serial Fragmentation
VSN: Variable Sorting for Normalization
MS-DAP: Mass Spectrometry Downstream Analysis Pipeline
FDR: False Detection Rate
SynGO: Synaptic Gene Ontology
ShinyGO: Shiny Gene Ontology
DIA: Data-Independent Acquisition
SNARE: Soluble NSF Attachment Receptor
GO: Gene Ontology
CHAPS: 3-[(3-cholamidopropyl)dimethylammonio]-1-propanesulfonate
WM: White Matter

---

**Table S1.**
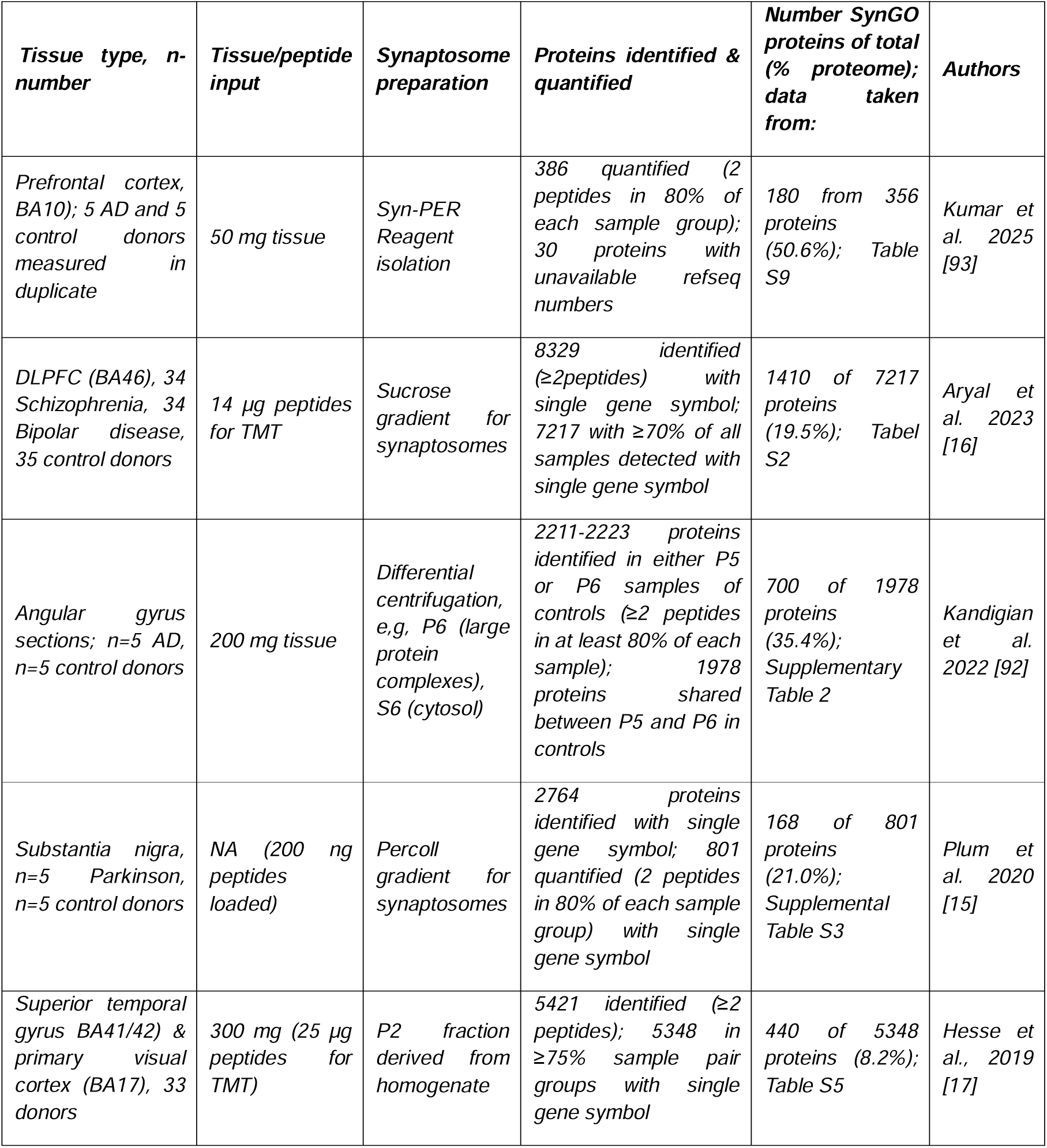
Overview of human proteomics studies focusing on the synapse, using similar cut-off for peptide quantification as used in the present study for comparison.

| *Tissue type, n-number* | *Tissue/peptide input* | *Synaptosome preparation* | *Proteins identified & quantified* | *Number SynGO proteins of total (% proteome); data taken from:* | *Authors* |
| --- | --- | --- | --- | --- | --- |
| *Prefrontal cortex, BA10); 5 AD and 5 control donors measured in duplicate* | *50 mg tissue* | *Syn-PER Reagent isolation* | *386 quantified (2 peptides in 80% of each sample group); 30 proteins with unavailable refseq numbers* | *180 from 356 proteins (50.6%); Table S9* | *Kumar et al. 2025 [93]* |
| *DLPFC (BA46), 34 Schizophrenia, 34 Bipolar disease, 35 control donors* | *14 µg peptides for TMT* | *Sucrose gradient for synaptosomes* | *8329 identified (≥2 peptides) with single gene symbol; 7217 with ≥70% of all samples detected with single gene symbol* | *1410 of 7217 proteins (19.5%); Tabel S2* | *Aryal et al. 2023 [16]* |
| *Angular gyrus sections; n=5 AD, n=5 control donors* | *200 mg tissue* | *Differential centrifugation, e.g. P6 (large protein complexes), S6 (cytosol)* | *2211-2223 proteins identified in either P5 or P6 samples of controls (≥2 peptides in at least 80% of each sample); 1978 proteins shared between P5 and P6 in controls* | *700 of 1978 proteins (35.4%); Supplementary Table 2* | *Kandigian et al. 2022 [92]* |
| *Substantia nigra, n=5 Parkinson, n=5 control donors* | *NA (200 ng peptides loaded)* | *Percoll gradient for synaptosomes* | *2764 proteins identified with single gene symbol; 801 quantified (2 peptides in 80% of each sample group) with single gene symbol* | *168 of 801 proteins (21.0%); Supplemental Table S3* | *Plum et al. 2020 [15]* |
| *Superior temporal gyrus BA41/42) & primary visual cortex (BA17), 33 donors* | *300 mg (25 µg peptides for TMT)* | *P2 fraction derived from homogenate* | *5421 identified (≥2 peptides); 5348 in ≥75% sample pair groups with single gene symbol* | *440 of 5348 proteins (8.2%); Table S5* | *Hesse et al., 2019 [17]* |

**Table S2:**
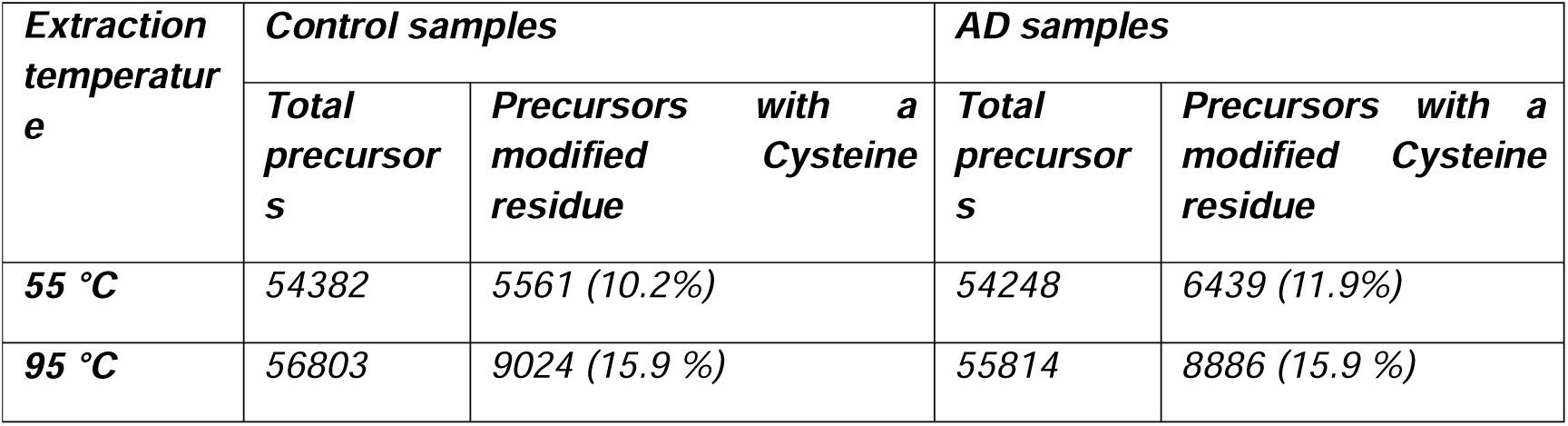
Impact of tissue extraction temperature on the efficiency of Cysteine reduction and alkylation. The average number of precursors identified consistently (75% of technical replicas) within the samples belonging to each incubation condition per donor type, both in total and when containing a modified (carbamidomethylated) cysteine residue.

| **Extraction temperature** | **Control samples** |  | **AD samples** |  |
| --- | --- | --- | --- | --- |
|  | **Total precursors** | **Precursors with a modified Cysteine residue** | **Total precursors** | **Precursors with a modified Cysteine residue** |
| **55 °C** | 54382 | 5561 (10.2%) | 54248 | 6439 (11.9%) |
| **95 °C** | 56803 | 9024 (15.9 %) | 55814 | 8886 (15.9 %) |

**Table S3.**
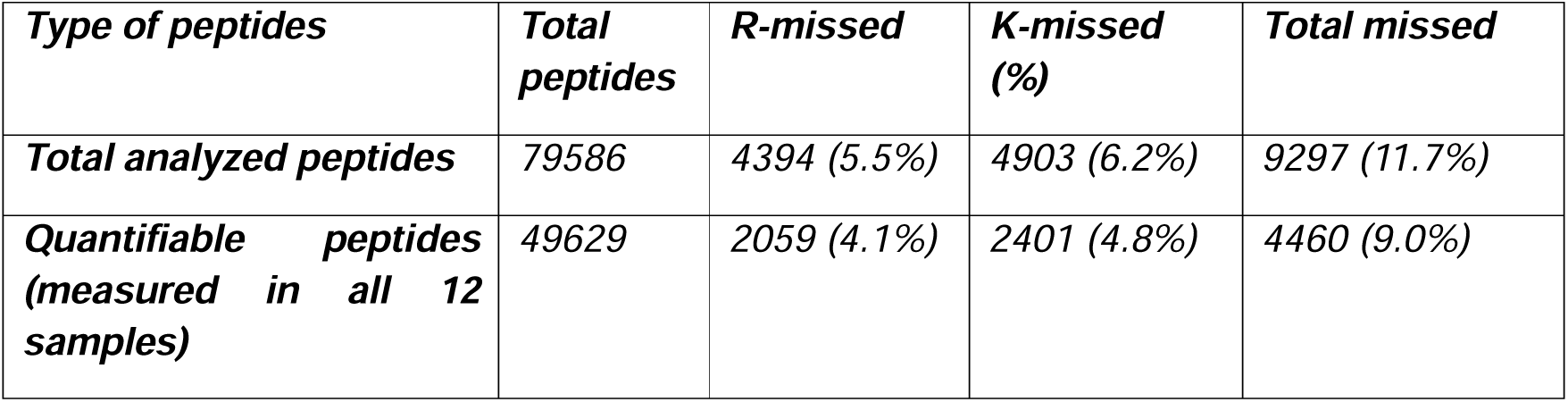
Number of peptides with missed Trypsin cleavage sites from Trypsin-pilot experiment. The number (%) of peptides with missed Arg (R) and Lys (K) sites from the total number of peptides analyzed (total peptides), as well as from the number of peptides that were present in all samples (i.e., quantifiable).

| **Type of peptides** | **Total peptides** | **R-missed** | **K-missed (%)** | **Total missed** |
| --- | --- | --- | --- | --- |
| **Total analyzed peptides** | 79586 | 4394 (5.5%) | 4903 (6.2%) | 9297 (11.7%) |
| **Quantifiable peptides (measured in all 12 samples)** | 49629 | 2059 (4.1%) | 2401 (4.8%) | 4460 (9.0%) |

**Figure S1.**
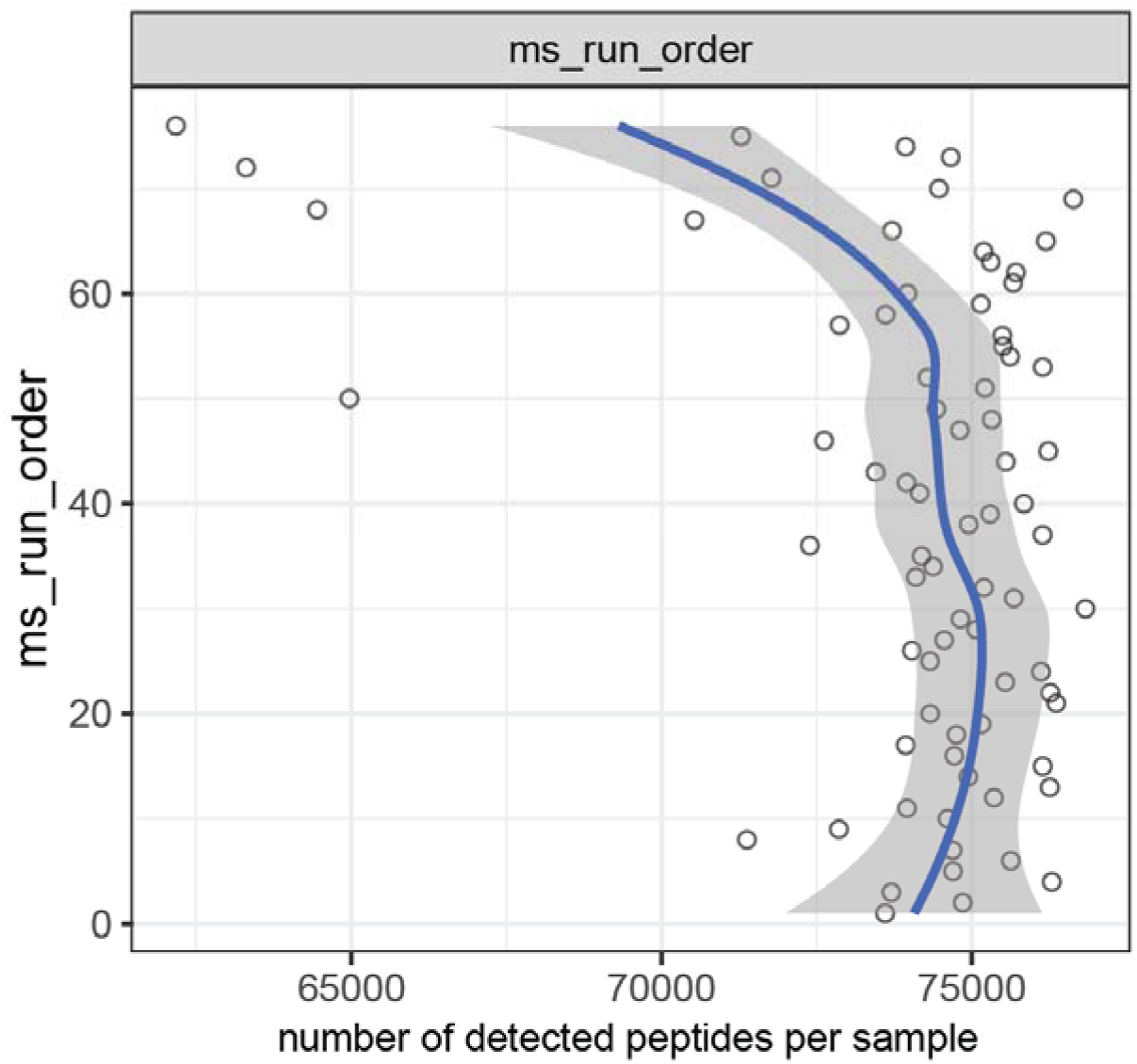
Necessity of randomization in MS workflow. One of the standard quality controls from the MS-DAP analysis shows the effect of the order in the MS analysis on the number of peptides detected. This emphasizes the importance of randomizing the samples across experimental groups to avoid technical effects biasing the results.

**Figure S2.**
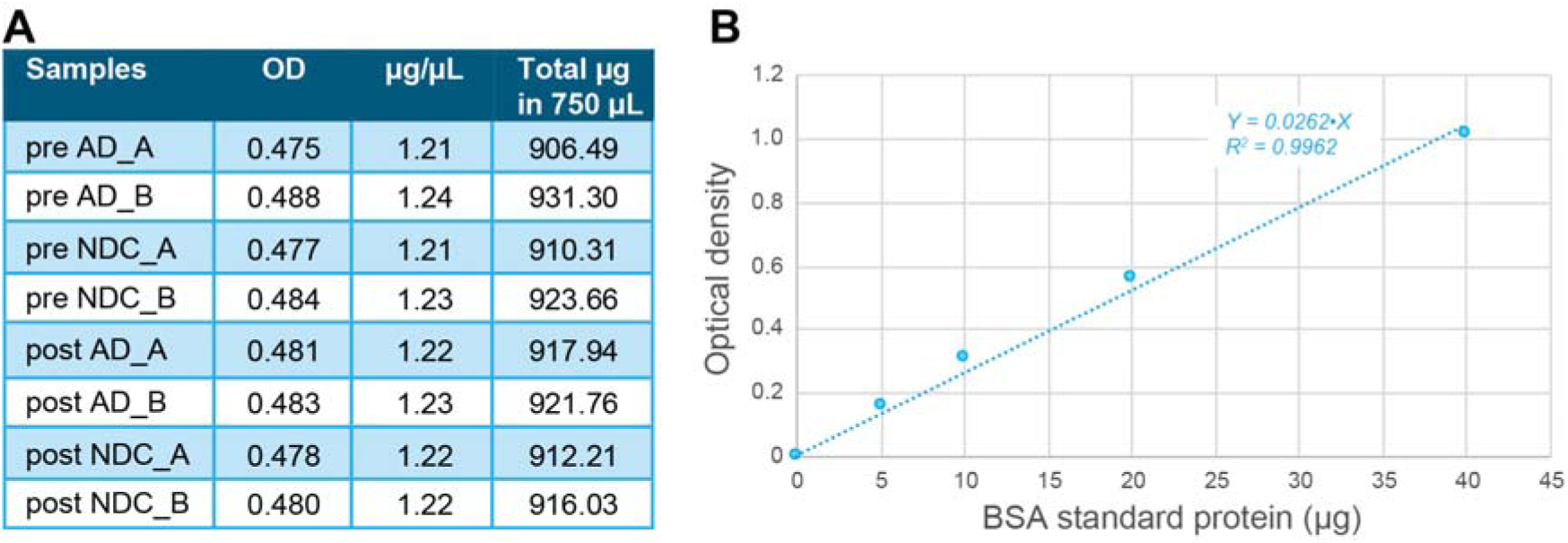
Determining isolated protein amount by BCA test. Optical density (OD) was measured for bovine serum albumin (BSA) standards (left) and was used to extrapolate protein concentration based on the OD of 15 µL from our samples (right). On average, our samples contained around 900 µg protein in 10 mg of initial tissue weight (dissolved in 750 µL 5% SDS for the OD measurements above). No weight loss was encountered by centrifugation. Pre = pre-centrifugation; Post = post-centrifugation; NDC = non-demented control donor; AD = donor with Alzheimer’s Disease.

**Figure S3.**
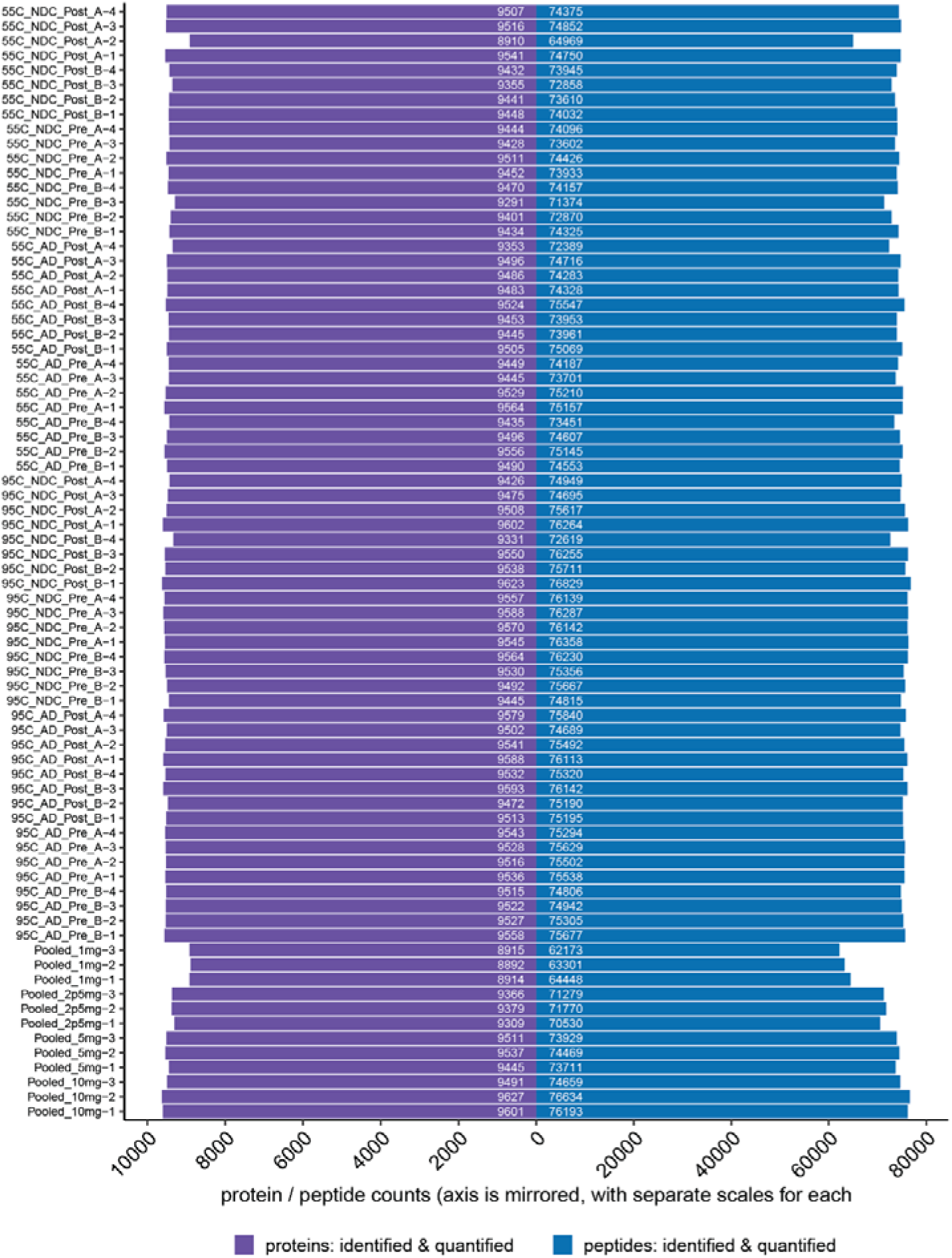
Number of peptides/proteins detected in pilot. This plot shows the number of (target) peptides that are detected per sample, as well as total protein number. For DIA, we refer to a peptide as ‘detected’ if the confidence score (for identification) is < 0.01.

**Figure S4.**
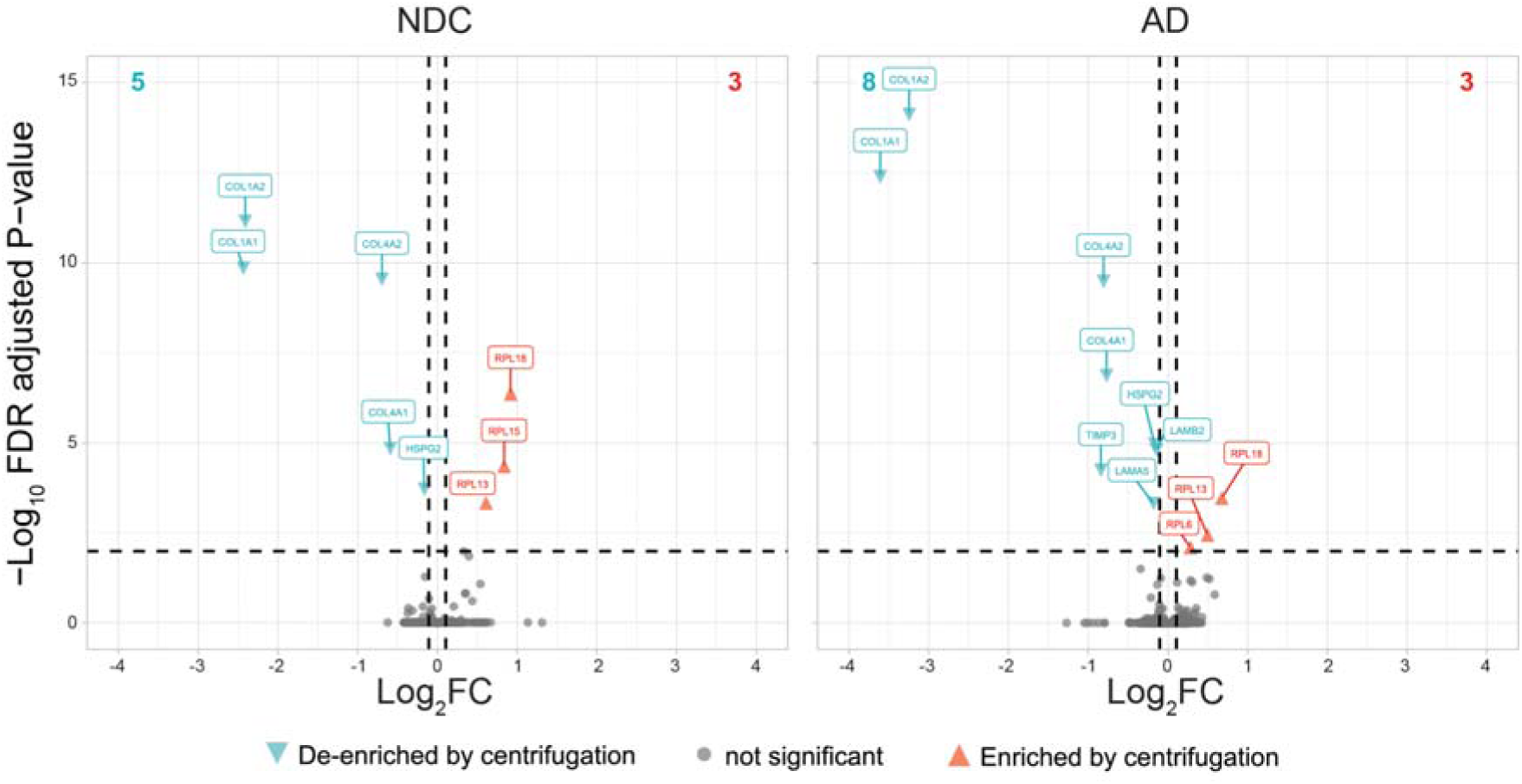
(De)enrichment of proteins by centrifugation is minimal. Volcano plots showing the DEqMS contrast of pre- vs. post-centrifugation for samples lysed at 55 °C, for control samples (NDC) and AD samples (AD). Gene symbols are shown for the significant proteins that are centrifugation-depleted (turquoise) or centrifugation-enriched (red) for adjusted p-values <0.01 (horizontal dashed line). The total number of differentially expressed proteins (colored; vertical dashed line; >|±0.10|) is indicated at the top (left, turquoise; right, red). Among the centrifugation-depleted proteins are multiple collagens. Proteins that are enriched by centrifugation include ribosomal proteins. Overall, the protein abundance decrease due to centrifugation is minimal, and greatly similar between the NDC and AD samples. The proteins for which no peptides were detected (truly “lost” proteins after centrifugation; not visible in this graph) were only a very small fraction of the total; Supplemental file 1.

**Figure S5.**
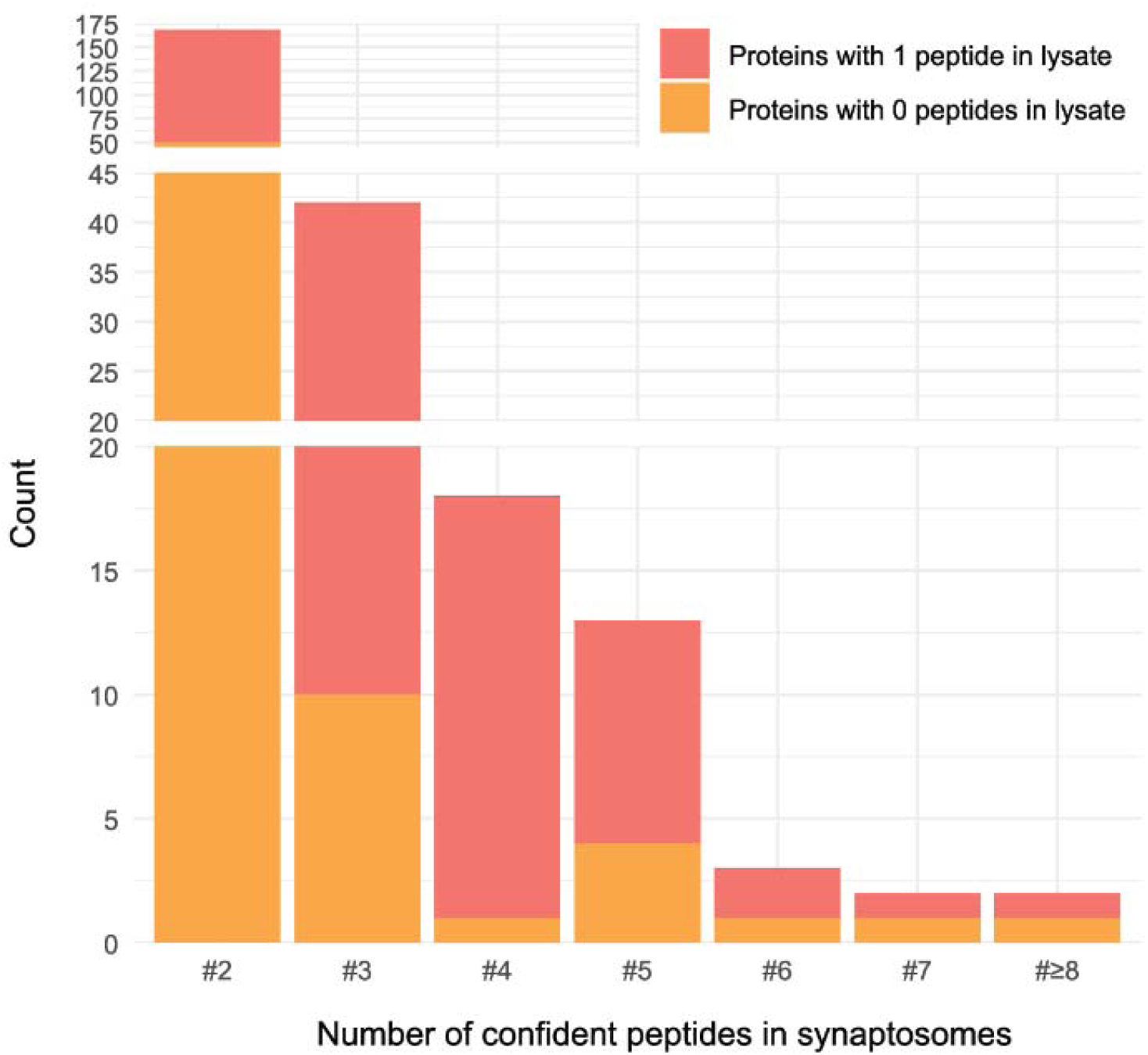
Proteins lost from lysate are often low-abundantly expressed and a small proportion is highly synaptosome enriched. Histogram of protein count detected in the lysate with 0 or 1 peptide (lost in lysate) by their peptide count detected in the synaptosome fraction. Proteins detected by 1 peptide in the lysate (n = 180) form the majority of these “lost” peptides and hence likely depend on stochastic events. Peptides confidently detected in synaptosomes from 4 and up (n = 38) form a group that are low expressed in lysate and highly enriched in synaptosomes.

**Figure S6.**
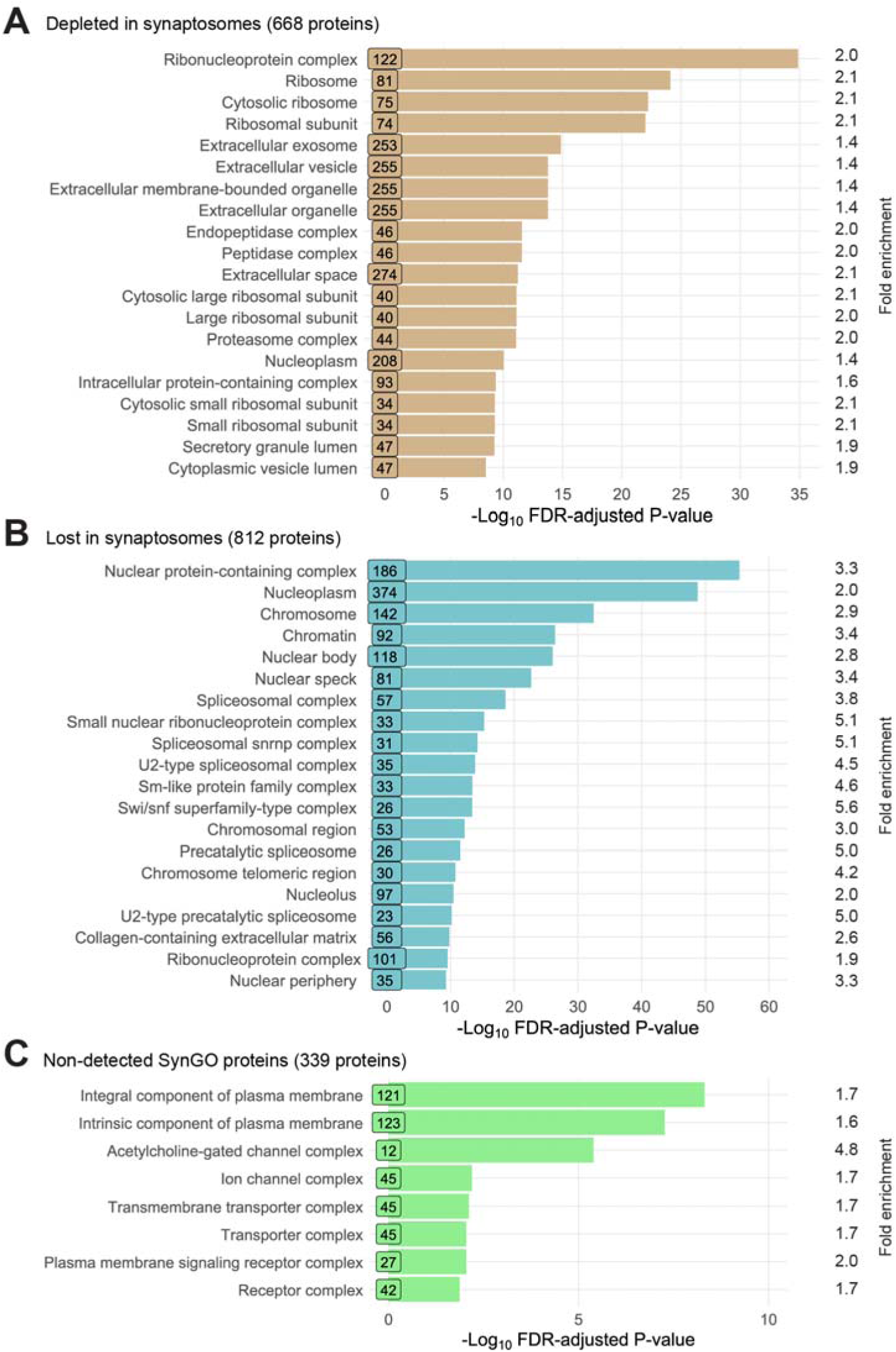
Protein pathways (cellular component, CC) of lost synaptosome proteins and absent SynGO proteins. A,B,C) Bar graphs (ShinyGO) showing the top 20 overrepresented CC-GO annotations of the 668 SynGO proteins that were depleted in the synaptosome isolation; *c.f.* Fig. 5A,E (A), the top 20 overrepresented CC-GO annotations of the 812 proteins that were not detected (“lost”) in the synaptosome isolation; *c.f.* Fig. 5A (B), and the 339 CC-SynGO proteins from the SynGO database that were not detected in either of the samples according to the criteria set; *c.f.* Fig. 5A (C). Note that the x-axis scale is different for panels A–C. For a complete list of proteins in each analysis, see Supplemental file 2.

**Figure S7.**
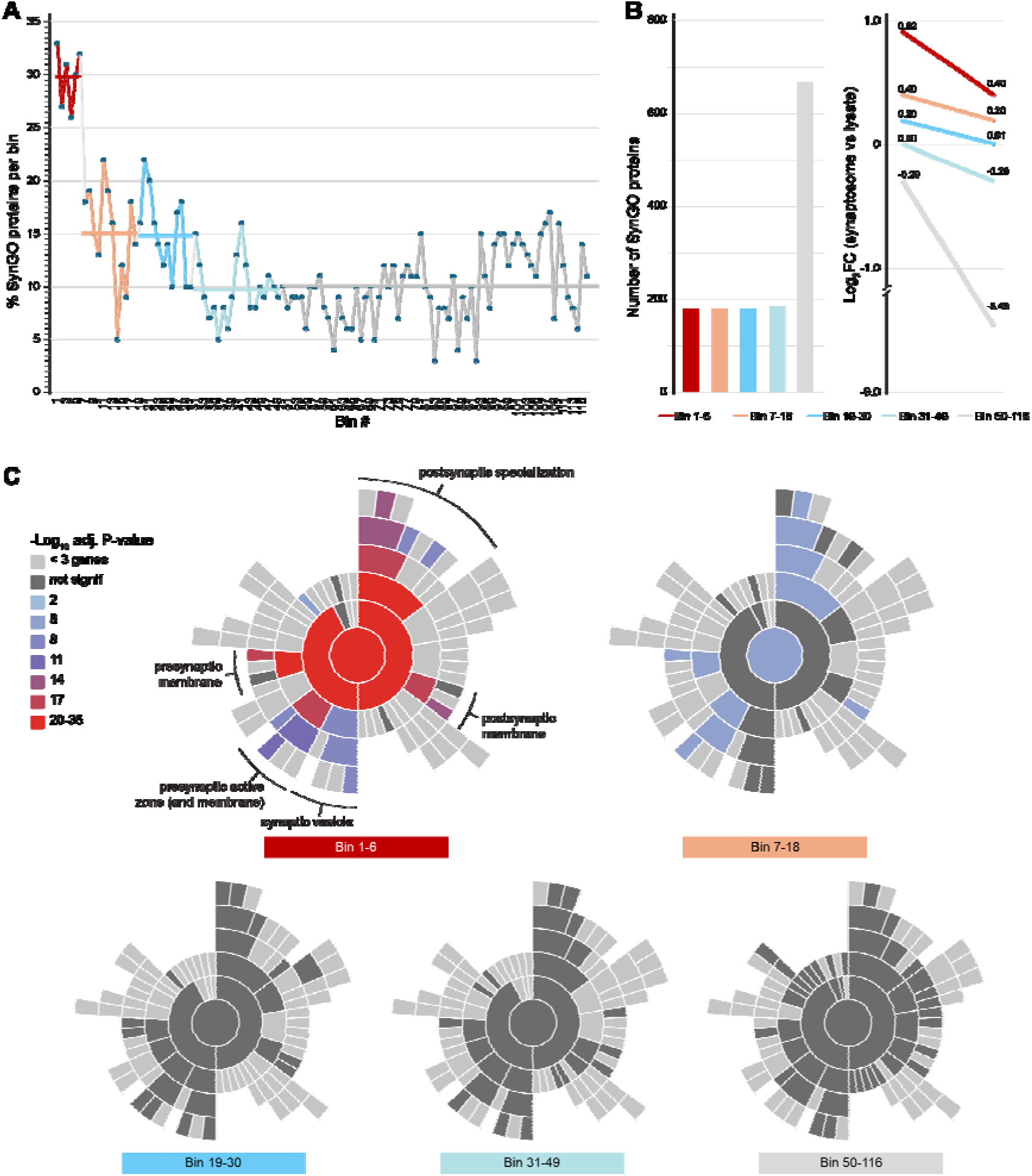
Binning of differential expression values indicates correct normalization between synaptosome and lysate samples. A) Plot showing the percentage of SynGO proteins among the 116 bins (50 proteins) based on differential expression between synaptosomes and whole-cell lysate samples (log2FC). Based on this, 5 main groups (color coded) were detected, the first showing the largest group of SynGO proteins. B) Bar graphs showing the total number of proteins per color-coded bin group (left) and the log2FC regulation values of each bin group (right). C) SynGO GSEA sunburst plots showing the overrepresentation analysis mainly in bins1-6 and 7-18 when compared to the total proteome detected.

**Figure S8.**
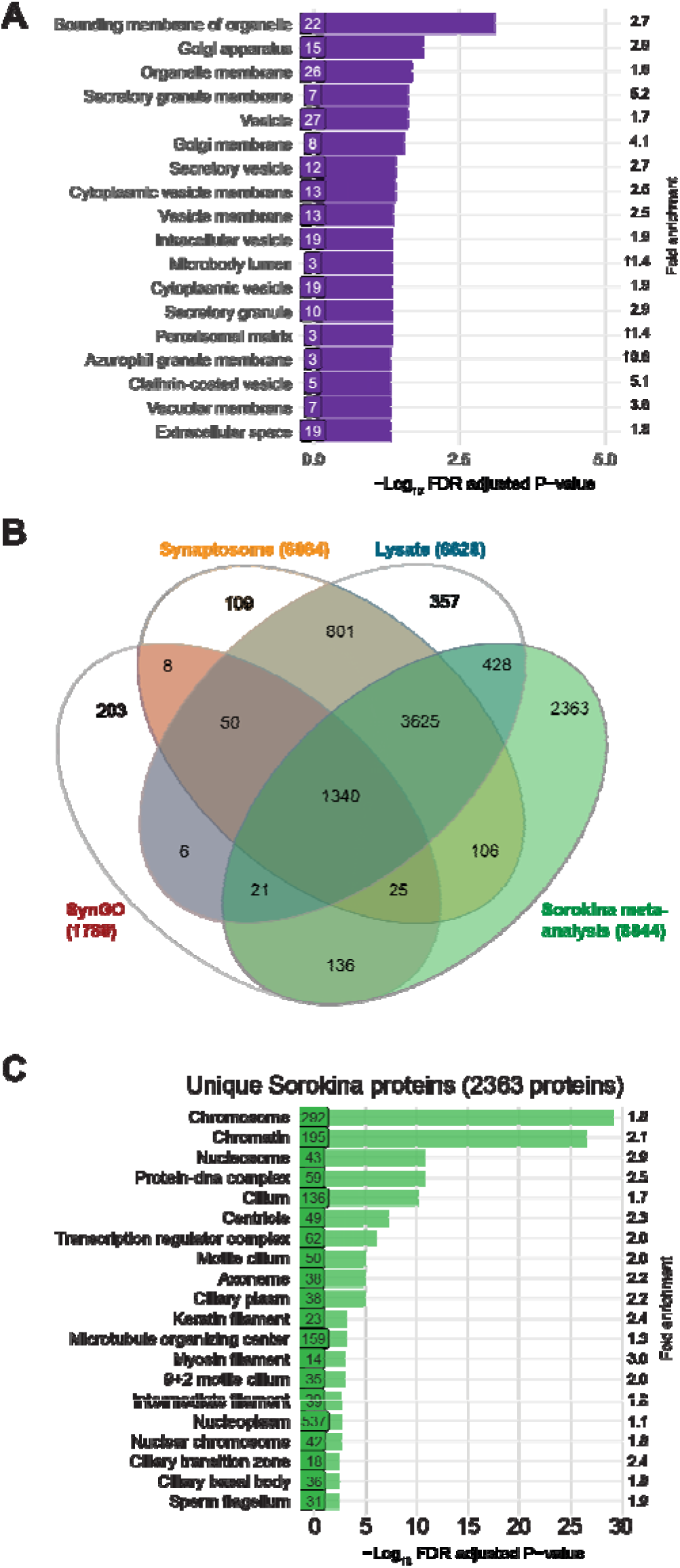
Protein pathways (cellular component, CC) of synaptosome-depleted human proteins and unique meta-data analysis synaptic proteins. A) Bar graph showing the 18 overrepresented cellular compartment CC-GO annotations (ShinyGO) of the 54 proteins that were enriched in synaptosomes in mouse [19] but were depleted in human synaptosomes. The number of proteins participating in each annotation, as well as the fold enrichment are indicated. B) Venn diagram showing the total number of detected proteins in synaptosome (“Synaptosome”), whole-tissue lysate (“Lysate”) samples (*c.f.* Fig 5A) and the 1789 SynGO annotated synaptic proteins (“SynGO”) compared with the 8044 unique proteins from the meta-analysis of the synaptic proteome as described by Sorokina et al. [67] . C) Bar graph showing the top 20 overrepresented CC-GO annotations of the 2363 proteins unique to the meta-analysis of the synaptic proteome [67] using ShinyGO. The number of proteins participating in each annotation, as well as the fold enrichment are indicated. For a complete list of proteins, see Supplemental file 2.

