## Supplemental Protocol 2_Synaptosome isolation for "A Tiered Approach to Human Synapse Proteomics: Optimized LC-MS/MS Analysis of Whole-Tissue and Synaptosome Preparations from Frozen Post-Mortem Brain Samples"

*Roig Kuhn, F., Klaassen, R.V., Koopmans, F.T.W., Koolman, T.S.Z., Smit, A.B., Spijker, S.*
*Dept. of Molecular and Cellular Neurobiology, Vrije Universiteit Amsterdam, Netherlands*

### Synaptosomes isolation protocol (small tissue samples)

This protocol uses a sucrose gradient on the S1 fraction of homogenized tissue. As input we typically use 25 mg human grey matter, sliced in 50 µm sections. When collecting the sectioned tissue, stack them on top of each other and 'fold' them to fit the tube. This has the advantage that it is easier to grind the tissue, because separate sections may float.

#### *Materials:*

#### Stock solutions

2 M Sucrose (MW 342.3) **0.5 L**

> Dissolve 342.3 g of sucrose in MilliQ (MQ) water by stirring. It might need to be heated in the shaker to completely dissolve. Fill up to 0.5 L after dissolving. Be sure that sucrose dissolves completely and there are no crystals in the bottle; after dissolving check the volume of the solution, if necessary adjust volume with MQ to 0.5 L. This solution can be stored at 4 °C and used during at least 4 months but check before every use that there is no fungi growth. Advice: store 45 mL aliquots in 50 mL Falcon tubes at -20 °C.

500 mM HEPES/NaOH pH 7.4 (MW 238.3) **50 mL**

> Check if the pH-meter has been calibrated properly with 2 standard solutions before any pH adjustments, if necessary, recalibrate it. Dissolve 5.96 g of HEPES in 40 mL of MQ water, adjust pH to 7.4 with freshly made 3 M NaOH (1.2 gram NaOH in 10 mL of MQ water), adjust volume to 50 mL.

Protease Inhibitor Cocktail Tablets, EDTA free

**Working concentration:** 1 tablet per 50 ml extraction solution
**Preparation of Working Solution:**

> One complete EDTA-free Protease Inhibitor Tablet is sufficient for the inhibition of the proteolytic activity in 50 ml extraction solution. When very high proteolytic activity is present, one tablet should be used for 25 ml extraction buffer. Tablets can be added directly to the extraction medium. Alternatively, a stock solution (25 X conc.) can be prepared.

**Preparation of stock solution (25 X conc.):**

> Dissolve one complete EDTA-free tablet in 2 mL dist. water or in 2 mL 100 mM phosphate buffer, pH 7.0.

**Storage conditions (working solution):**

> The stock solution can be stored at 2 to 8 °C for 1 to 2 weeks, or at least 12 weeks at -15 to -25 °C. Use the same batch of solutions for a complete experiment.

*Roig Kuhn, F., Klaassen, R.V., Koopmans, F.T.W., Koolman, T.S.Z., Smit, A.B., Spijker, S.*
*Dept. of Molecular and Cellular Neurobiology, Vrije Universiteit Amsterdam, Netherlands*

#### **Working buffers** (make at least one day prior to use)

A. **Homogenisation Buffer:**
   5 mM HEPES/NaOH pH 7.4
   0.32 M Sucrose
B. **0.85 M Sucrose:**
   5 mM HEPES/NaOH pH 7.4
   0.85 M Sucrose
C. **1.2 M Sucrose:**
   5 mM HEPES/NaOH pH 7.4
   1.2 M Sucrose
D. **5 mM HEPES:**
   5 mM HEPES/NaOH pH 7.4

| Buffer | 50 mL | | | 300 mL | | |
|---|---|---|---|---|---|---|
| | 500 mM HEPES (in mL) | 2 M Sucrose (in mL) | H2O (in mL) | 500 mM HEPES (in mL) | 2 M Sucrose (in mL) | H2O (in mL) |
| Homogenisation | 0.5 | 8 | 41.5 | 3 | 48 | 249 |
| 0.85 M Sucrose | 0.5 | 21.25 | 28.25 | 3 | 127.5 | 169.5 |
| 1.2 M Sucrose | 0.5 | 30 | 19.5 | 3 | 180 | 117 |
| 5 mM HEPES | 0.5 | - | 49.5 | 3 | - | 297 |

50 mL buffers can be made in blue capped Falcon tubes. After preparation keep them at 4 °C or store at -20 °C for long-term use.

Check clean glassware availability

*Roig Kuhn, F., Klaassen, R.V., Koopmans, F.T.W., Koolman, T.S.Z., Smit, A.B., Spijker, S.*
*Dept. of Molecular and Cellular Neurobiology, Vrije Universiteit Amsterdam, Netherlands*

### *Methods:*

##### *Prior to the isolation day*

1. Put the rotor and buckets for the ultracentrifuge at 4 °C to save cooling time the next day

##### *The isolation day*

First things to do:

1. Switch on the low-speed Eppendorf centrifuge 5810R (with rotor for 50 mL Falcon tubes) and run the *"Fast cooling"* program
2. Switch on the ultracentrifuge, make sure the correct rotor and tubes are set. Set the correct speed/time/temperature. Put the rotor in, close the lid and press vacuum
3. Take ice and put the potter homogeniser (5 mL potter tube) on ice with the Teflon pestle inside
4. Put labelled 15 mL Falcon tubes on ice, one for every sample
5. Prepare the homogenisation buffer with inhibitors (for 6 samples):
   In a 50 mL tube: 50 mL of the homogenisation buffer, add 1 tablet of the Complete <u>EDTA-free</u> Roche inhibitory cocktail. Put on the roller bank until the tablet is completely dissolved. Put on ice
6. Take a pair of tweezers (pincers; sharp ends): not all of them can reach the bottom of a 1.5 mL Eppendorf tube. Make sure the tweezers are cold
7. When ready and the homogenisation buffer is cold, **<u>start</u>**:

##### I. Tissue Homogenisation

8. Attach the cold Teflon pestle to the potter and set the homogenizer to 900 rpm (do not switch on yet); Fill the beaker with 50% normal ice and 50% water to create a large contact area for the tube. Make sure the pestle reaches the bottom of the tube
9. Ready the homogeniser: fixate the pestle into place and leave potter tube on ice;
10. Take out the 1.5 mL tube with brain tissue from -80 °C or from box of dry ice. Add 1 mL of homogenisation buffer to the tube. Cut off about 1 mm from the tip of a P1000 pipet, and mix the tissue sample with the homogenisation buffer by pipetting up and down, then vortex briefly until the tissue is suspended. Transfer the sample in the buffer to the potter tube (on ice). Add another 1 mL homogenisation buffer to the 1.5 mL tube and repeat the previous steps, then transfer into the potter for a total volume of 2 mL sample
11. Place the Teflon pestle into the Potter tube with your sample, ***and only after that!!!*** switch on the homogenizer – at **900 rpm**;
12. Homogenize the sample with **12 slow strokes at 900 rpm (moving all the way up and down without taking the pestle out of the buffer)**; (if necessary: take an aliquot homogenate sample and store at -80 °C)
13. Carefully transfer the homogenate to the labelled tube (15 mL Falcon tube) on ice;
14. Add 3.5 mL homogenisation buffer for a total of ~5.5 mL sample volume to the 2 mL homogenate
15. Wash the Potter and its pestle well with soap and water, then rinse with MQ and put on ice (with the pestle) for 2-3 min

16. When cooled down start with the next sample

**II. Low speed centrifugation: purification from cell debris**

17. Centrifuge **1,000x g at 4 °C** for 10 min. Keep the time between centrifugation and loading on the sucrose gradient as short as possible. If necessary, centrifuge after sucrose gradient is prepared (step 19-23)
18. For taking off the supernatant of this step, see step 24

**III. Ultracentrifugation: 0.85–1.2M sucrose step gradient:**

19. While the samples are being centrifuged, establish a step gradient in tubes for ultracentrifugation:

| Sample volume (in mL) | Centrifuge buckets | Sucrose buffers | Sample volume |
|---|---|---|---|
| ≤ 5 mL | 'small' (16 mL total) | 5 mL 1.2 M (C)<br>5 mL 0.85 M (B) | 5.0–5.5 mL |
| > 5 – 17 mL | 'big' (37 mL total) | 12 mL 1.2 M (C)<br>12 mL 0.85 M (B) | 15-17 mL |

20. Use ~17 mL ultracentrifugation tubes. The sample volume is ~5 mL so:
21. Pipet (volume pipette) 5 mL 1.2 M sucrose into tube
22. Pipet 5 mL 0.85 M sucrose on top to get two clear phases. Note: pipet <u>carefully and very slowly</u>, while keeping tube under an **<u>angle</u>**!
23. After making the gradient, put tube on ice
24. Take tubes with sample from low-speed centrifuge and put them on ice (see II, step 17)
25. When transferring the supernatant (S1 fraction) try <u>not to touch the bottom of the tube</u> with your pipette tip. Very carefully put the supernatant <u>on top</u> of the sucrose step gradient (~5.5 mL)
26. Be sure that a maximum of 5-7 mm space is left between the surface level of the solution and the edges of the tube (if not - adjust the level with the Homogenisation buffer). Otherwise edges of the tube <u>may collapse</u> in the ultracentrifuge!
27. Balance weights of buckets **with** tubes on a balance using homogenisation buffer. The difference between two opposite tubes (1-4; 2-5; 3-6) should not exceed 0.02 gram!! When using 4 tubes, leave the two remaining (coupled = opposite!) buckets empty, but **do put them in the centrifuge**
28. Centrifuge at 32,000 rpm at 4 °C for 2 hours
29. Collect the white disk of the 1.2 – 0.85 M interphase (=synaptosomes)
    <u>Advice</u>: First use a plastic Pasteur pipette to take off from the top all that you don't want (leave a bit). Then take a fresh pipette (or use a yellow tip on a blue 1 mL tip on the pipet; set at 750 µL) to carefully collect the white disk at the interphase and transfer to a fresh 5 mL Eppendorf tube

*Roig Kuhn, F., Klaassen, R.V., Koopmans, F.T.W., Koolman, T.S.Z., Smit, A.B., Spijker, S.*
*Dept. of Molecular and Cellular Neurobiology, Vrije Universiteit Amsterdam, Netherlands*

30. To the collected sample (~ 1.5 mL), add ~ 3.0 mL 5 mM HEPES in a 5 mL Eppendorf tube, immediately invert a few times (end concentration sucrose ~0.32 M). Centrifuge the samples at 18,000x g (Eppendorf 5810) at 4 °C for 30 minutes
31. Remove supernatant carefully with pipet (watch out, the pellet is loose)
32. Resuspend pellet in 75 µL 25 mM homogenisation buffer (volume depends on the starting material). Keep 5 µl for Bradford assay (if performing step 33), and aliquot (2 x 35 µL) the rest for synaptosome proteomics (see step 1 of '<u>Synaptosome protein extraction</u>'). Store these samples at -80 °C.
33. Measure the concentration with Bradford to have a first assessment of the protein yield (in case you proceed with other techniques than LC-MS)
