## Supplemental Protocol 1_Pre-MS workflow for "A Tiered Approach to Human Synapse Proteomics: Optimized LC-MS/MS Analysis of Whole-Tissue and Synaptosome Preparations from Frozen Post-Mortem Brain Samples"

*<u>Supplemental protocol 1</u>: **Detailed pre-MS workflow protocol** accompanying "An optimized quantitative MS-based neuroproteomics approach for frozen post-mortem brain samples"*

*Roig Kuhn, F., Klaassen, R.V., Koopmans, F.T.W., Koolman, T.S.Z., Smit, A.B., Spijker, S.*
*Dept. of Molecular and Cellular Neurobiology, Vrije Universiteit Amsterdam, Netherlands*

### Detailed pre-MS workflow protocol

➢ Schematic overview of the **<u>pre-MS workflow:</u>**

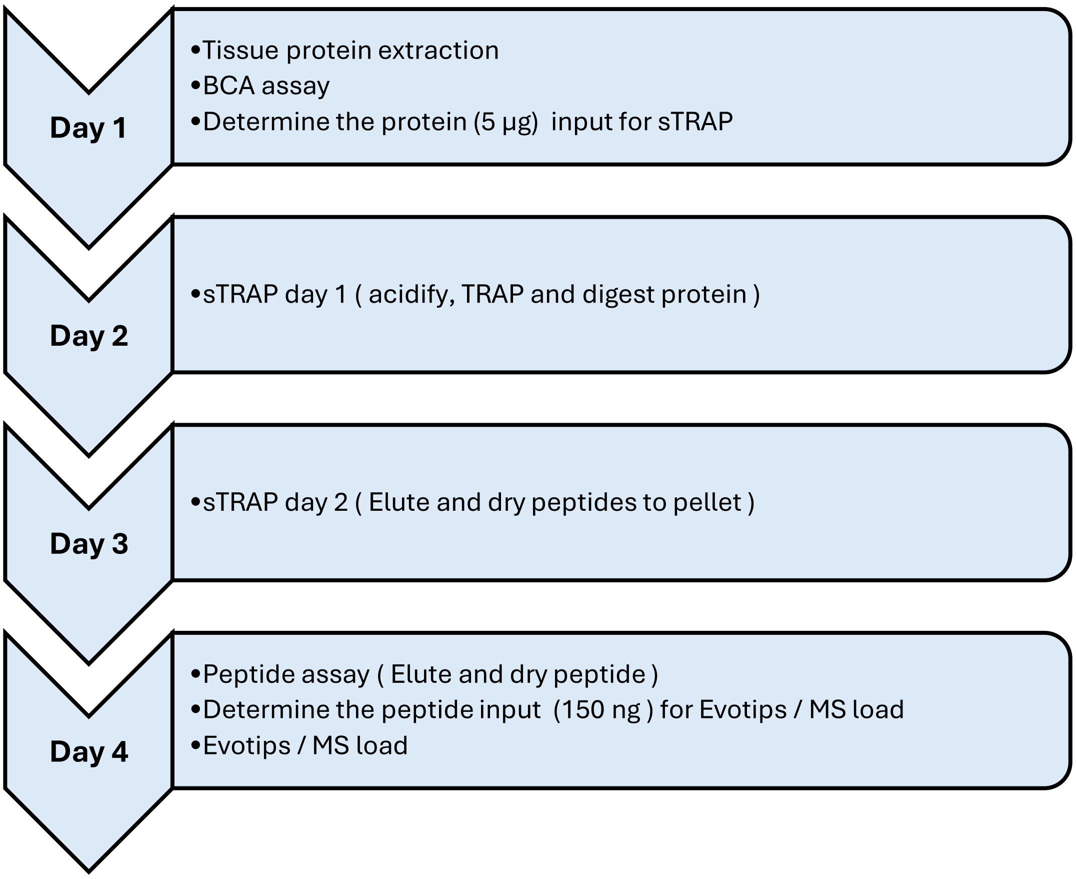

*Roig Kuhn, F., Klaassen, R.V., Koopmans, F.T.W., Koolman, T.S.Z., Smit, A.B., Spijker, S.*
*Dept. of Molecular and Cellular Neurobiology, Vrije Universiteit Amsterdam, Netherlands*

#### Day 1: Protein extraction

##### Material:

- Stocks:
  20% SDS (Sodium Dodecyl Sulfate)
  1 M Tris-HCl buffer pH 8.0
  100 mM TCEP (Tris(2-CarboxyEthyl)Phosphine)
  200 mM 2-CAA (2-ChloroAcetamide)
- 1.5 mL Snap-lock Eppendorf tube
- Thermomixer, centrifuge, dry ice, and wet ice

##### Preparation:

- Warm up the thermomixer to 95 °C
- Fast cool the centrifuge to 4 °C
- Make **40 mL** of 2x extraction buffer (10% SDS, 100 mM Tris-HCl buffer pH 8.0, 10 mM TCEP, 40 mM 2-CAA)

| **2x Extraction buffer: 10% SDS, 100 mM Tris-HCl buffer pH 8.0, 10 mM TCEP, 40 mM 2-CAA** | | | |
|---|---|---|---|
| Material | Stock concentration | Required concentration | Volume from stock ( mL ) |
| MQ-water | - | - | 4 |
| Tris-HCl buffer pH 8.0 (M) | 1 | 0,1 | 4 |
| TCEP (mM) | 100 | 10 | 4 |
| 2-CAA (mM) | 200 | 40 | 8 |
| SDS (%) | 20 | 10 | 20 |
| End volume | | | 40 |

- Make **40 mL** of 1x extraction buffer by diluting (1:1) the 2x extraction buffer with MQ-water

##### Method:

##### Tissue protein extraction

1. Transport the sample from the -80 °C freezer to the lab on dry ice.
2. Centrifuge the samples: 10,000 x g, 4 °C for 10 seconds to pellet the tissue to the bottom of the tube.
3. Place the sample on wet ice and let it thaw for 2 minutes.
4. Add **500 µL** of 1x extraction buffer to the sample and vortex: 2,200 rpm at RT until the tissue is fully dissolved. Vortex on the cap briefly if needed to ensure complete extraction.
5. Incubate the sample in the thermomixer for 15 minutes at 95 °C and 1500 rpm. Let the sample cool in the same thermomixer for 15 minutes at 55 °C and 1500 rpm.
6. After 30 minutes of total incubation time, remove the sample from the thermomixer and allow it to cool to room temperature.
7. Spin the sample down in the centrifuge: 10,000 x g (rcf) at RT for 5 minutes.
8. Aliquot the sample for BCA assay ***(7.5 µL & 15 µL)*** and label on the tube the aliquoted sample volume.
9. Add **7.5 µL** 1x extraction buffer to the 7.5 µL BCA sample. ***(end volume = 15 µL)***
10. Aliquot ***40 µL*** sample for STRAP and freeze it at -80 °C along with the remaining sample. ***(437.5 µL)***

##### Synaptosome protein extraction

1. Transport the extracted synaptosome sample (see Supplemental protocol 2 for synaptosome isolation) from the -80 °C freezer on wet ice.
2. Let the sample slowly thaw to room temperature
3. Vortex sample briefly: 2, 000 rpm at RT and place sample back on ice.

*Roig Kuhn, F., Klaassen, R.V., Koopmans, F.T.W., Koolman, T.S.Z., Smit, A.B., Spijker, S.*
*Dept. of Molecular and Cellular Neurobiology, Vrije Universiteit Amsterdam, Netherlands*

4. Centrifuge the samples: 10,000 x g at 4°C for 30 seconds to collect all fluid to the bottom of the tube.
5. Transfer **35 µL** of the synaptosome to a new 1.5 mL Eppendorf tube.
6. Add **35 µL** of 2x extraction buffer to the sample ***(end volume sample = 70 µL.)***
7. Incubate the sample in the thermomixer for 15 minutes at 95 °C and 1500 rpm. Let the sample cool in the same thermomixer for 15 minutes at 55 °C and 1500 rpm.
8. After 30 minutes of total incubation time, remove the sample from the thermomixer and allow it to cool to room temperature
9. Centrifuge the sample: 10,000 x g at RT for 5 minutes.
10. Aliquot the sample for BCA assay ***(7.5 µL & 15 µL)*** and label appropriately.
11. Add **7.5 µL** 1x extraction buffer to the 7.5 µL BCA sample. ***(end volume = 15 µL)***
12. Aliquot ***40 µL*** sample for STRAP and freeze it at -80°C along with the remaining sample. ***(7.5 µL)***

#### BCA protein Assay

##### Material:

- 2x extraction buffer (10% SDS, 100 mM Tris-HCl buffer pH 8.0, 10 mM TCEP, 40 mM 2-CAA)
- 1x extraction buffer (5% SDS, 50 mM Tris-HCl buffer pH 8.0, 5 mM TCEP, 20 mM 2-CAA)
- BCA protein assay kit (Pierce, Thermo Scientific - 23225)
  96-well plate, clear bottom
- 1.5 mL Safe-lock Eppendorf tube
- Microplate reader (SpectraMax, Molecular Devices)

##### Preparation:

- Prepare BSA standard dilutions (15.0 – 12.0 – 10.0 – 7.5 – 5.0 – 2.5 – 1.25 – 0 µg/µL) and aliquot **15 µL** of each standard in replica tube. For the blank, use **15µL** of 1x extraction buffer

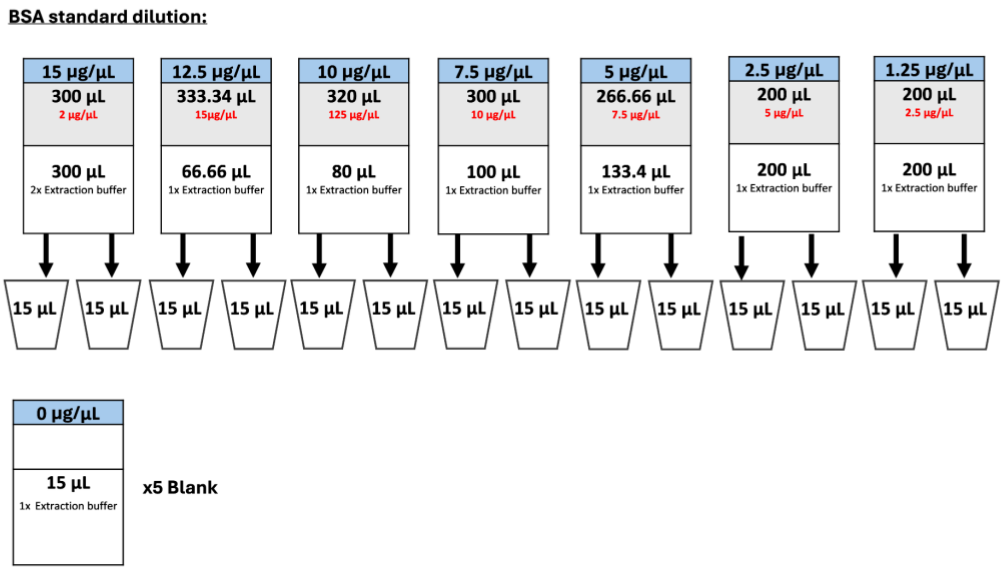


- Make BCA working reagent A:B (1:50) from the BCA protein assay kit for the required number of samples. This includes:
  a. 21 BSA standard curve samples (8 concentrations in duplicate + 5 blanks)
  b. Number of original samples with **each**:

i. 2x 15 µL from step 8/9 (tissue protein extraction) **or**
ii. 2x 15 µL from step 10/11 (synaptosome protein extraction)

For every BCA analysis, 200 µL BCA working reagent is used, hence making the final volume for at least 5 samples extra (18 mL –see Table below– is sufficient for 32 original samples (measured in duplicate) in addition to the 21 BSA standard curve samples and 5 spare reactions)

| **Working reagents A:B (1:50)** | |
|---|---|
| **Material** | **Volume (mL)** |
| Solution A | 17.6 |
| Solution B | 0.4 |
| **End volume** | **18** |

- Take the aliquot of extracted protein samples for BCA assay (the labeled '7.5' and '15' from step 8/9 (tissue protein extraction) or step 11/12 (synaptosome protein extraction) labeled '7.5' and '15').
- Determine the 96-well plate layout for the assay.

#### Method:

1. Mix the extracted protein samples (labeled '7.5' and '15') in the thermomixer: 1,450 rpm at 37 °C for 2 minutes.
2. Centrifuge all samples (extracted protein samples, BSA, and blanks): 10,000 x g at RT for 10 seconds to collect all liquid to the bottom of the tube.
3. Add **200 µL** of BCA working reagent to all samples and blanks. Use a repeating pipette and note down the start time. ***(end volume = 215 µL)***
4. Mix the sample by gently inverting the tubes.
5. Centrifuge the samples: 10,000 x g at RT for 10 seconds to collect all fluid to the bottom of the tube.
6. Transfer **200 µL** of the combined sample and BCA working reagent to the 96-well plate. Avoid bubbles in the plate during pipetting, this will interfere with the absorption reading.
7. Incubate the plate at room temperature for 30 minutes (from the start time of step 3), 45 minutes, and 60 minutes.
8. Measure absorbance at each time point (30, 45, and 60 minutes) using a microplate reader.

#### Settings for the microplate reader (SpectraMax):

**Optical configuration:** Monochromator
**Read mode:** Absorption
**Read Type**: Endpoint
**Emission lm1:** 562nm
**Plate type >**
**Plate format:** 96 wells
**Select specific:** 96 well plate standard crlbtm

#### BSA standard curve

1. Determine the average absorption for the blanks.
2. Determine the average absorption value for each standard.
3. Subtract the average blank from average absorption value for each standard.

*Absorption value standard – average blank = corrected absorption value of standard*

4. Insert a scatter chart using the value from the corrected absorption BSA. Insert a trendline, set the intercept to 0, and display the equation and the $R^2$ on the chart.

#### Amount of protein in the sample

5. Determine the average absorption for each sample
6. Correct the average absorption for each sample.
7. 

*Absorption value sample – average blank = corrected absorption value sample*

8. Determine the amount of protein (µg) in the isolated protein samples.

*Corrected absorption value sample / linear slope = protein in 7.5 µL or 15 µL*

9. Determine the amount of protein (µg) in 1 µL (divide by 7.5 for the '7.5' sample and by 15 for the '15' µL sample)

#### Protein volume for STRAP (10µg in 50µL)

10. Calculate for each sample the amount of volume needed for 10µg protein.

*10 (µg) / the amount of protein (µg) in 1 µL = volume needed for 10 µg of protein*

11. Calculate the required volume for each sample to achieve an end volume of 50 µL sample. 10µg in 50µL for STRAP.

*50 (µL) - volume for 10 µg of protein = volume 1x extraction buffer*

**This is the end of day 1**<u>. Please proceed to page 6 for the sTRAP to be performed on day 2 & 3.</u>

#### Day 2: sTRAP day 1

##### Material:

- 1x extraction buffer
- Ammonium bicarbonate (ABC), methanol, 85% phosphoric acid, 1 M Tris-HCl buffer pH 8.0
- Thermomixer, centrifuge, wet ice
- HiPure DNA Micro column (Magen - C13011), 1.5 mL protein Lobind / Safe-lock Eppendorf tube.
- Trypsin Lys-C vial 20 µg (Promega - 254661-1)
- Humid chamber/box filled with tissue soaked in water

##### Preparation:

- Make **10 mL** of 1 M Ammonium bicarbonate in a 15 mL tube (ABC MW: 79.056 g/mol)

| 1 M Ammonium bicarbonate (ABC) in MQ-water | |
|---|---|
| Material | Amount |
| ABC | 0.79 gram |
| MQ-water | Fill until 10 mL |

- Make **40 mL** of 50 mM ABC in a clean glass bottle

| 50 mM Ammonium bicarbonate (ABC) in MQ-water | | | |
|---|---|---|---|
| Material | Stock concentration | Required concentration | Volume from stock (mL) |
| ABC (M) | 1 | 0.05 | 2 |
| MQ-water | - | - | 38 |
| End volume | | | 40 |

- Make **40 mL** of 12% phosphoric acid in a clean glass bottle.

| **Acidifier: 12%** phosphoric acid in MQ- water | | | |
|---|---|---|---|
| Material | Stock concentration | Required concentration | Volume (mL) |
| Phosphoric acid (mL) | 85 | 12 | 5.6 |
| MQ-water (mL) | - | - | 34.4 |
| End volume | | | 40 |

- Make **150 mL** of binding/wash buffer that contains (9:1) methanol and 100 mM Tris-HCl buffer pH 8.0

| **Binding/wash buffer: (9:1) 90%** methanol diluted in **100 mM** Tris buffer pH 8.0 | | | |
|---|---|---|---|
| Material | Stock concentration | Required concentration | Volume (mL) |
| Tris-HCL buffer pH 8.0 (M) | 1 | 0.1 | 15 |
| Methanol (%) | 100 | 90 | 135 |
| End volume | | | 150 |

- Determine the required amount of digestion buffer (trypsin/Lys-C in 50 mM ABC) for each sample. Each sample should receive a 1:25 ratio of protein weight to trypsin weight in 50 µL of digestion buffer.
  *For example, for 10 µg of protein to digest, apply 0.4 µg trypsin/Lys-C in 50 mM ABC to each sample in an end volume of 50 µL*
- Label tubes/column

#### Method:

#### 10 µg of protein per sample

1. Thaw the sTRAP sample (from day 1, **step 10/12** tissue/synaptosomes) by transferring it from the -80 °C freezer and allowing it to thaw at room temperature.
2. Centrifuge the sample: 20,000 x g at RT for 10 seconds to collect all fluid to the bottom of the tube.
3. Warm the sample in the thermomixer: 1,400 rpm at 37 °C for 2 minutes to redissolve the SDS back in solution.
4. Centrifuge sample: 20,000 x g for 5 minutes at RT to pellet any debris.

5. Add 1x extraction buffer volume * for each sample in a new 1.5 mL Safe-lock Eppendorf tube.
   **See the BCA assay results for the exact volume per sample.*
6. Pipette the correct amount of extracted protein for each sample * to the 1x extraction buffer:
   **See the BCA assay results for the exact volume per sample.*

   **volume 1x extraction buffer + volume extracted protein = end volume 50 µL sample**

7. Mix the sample in the thermomixer: 1,400 rpm at RT for 5 minutes.
8. Centrifuge the sample: 10,000 x g at RT for 1 minute to collect all fluid to the bottom of the tube.

##### Acidify and TRAP protein

9. Add **5 µL** of 12% phosphoric acid to the 50 µL samples of step 6. The final concentration of **phosphoric acid = 1.1% (1:10 dilution)**
10. Vortex the sample briefly: 1,500 rpm at RT.
11. Add **350 µL** of binding/wash buffer to the sample mix.
12. Vortex the sample briefly: 1,500 rpm at RT.
13. Transfer **450 µL** (the entire sample) to the column placed in a flow-through collector.
14. Centrifuge the column: 1,450 x g at RT for 1 minute. The protein will now be trapped in the column.
    *Visually confirm that all of the samples have passed through the column. If not, centrifuge again.*

#### Wash/clean protein and make the trypsin-Lys-C digestion buffer.

15. Wash the column with **400 µL** binding/wash buffer
16. Centrifuge the column: 1,450 x g at RT for 1 minute. Turn the column 180 degrees between centrifugation steps (see below) to ensure an even wash of the column.

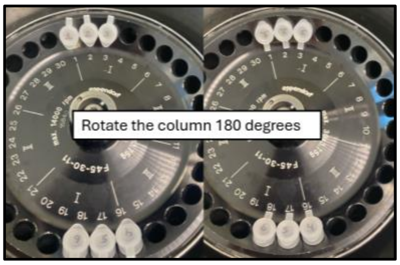


17. Discard the flow through and repeat steps 15 until 17 a total of four times. Be sure to turn the column between each centrifugation step. **Do not let the columns dry in between washes**.
    ***Prepare the digestion buffer during the fourth wash step to prevent the column from drying.***

*Roig Kuhn, F., Klaassen, R.V., Koopmans, F.T.W., Koolman, T.S.Z., Smit, A.B., Spijker, S.*
*Dept. of Molecular and Cellular Neurobiology, Vrije Universiteit Amsterdam, Netherlands*

***Add 400 µL binding/wash buffer and do not proceed with centrifugation (step 17) until the digestion buffer is ready. Use 50 mM ABC buffer to dissolve the trypsin and keep it on ice.***

18. Centrifuge the column: 1,450 x g at RT for 1 minute. Inspect the column to ensure that all the binding/wash buffer has passed through. If necessary, centrifuge again. If the binding/wash buffer is not completely removed, the trypsin may elute from the column before or during the overnight incubation.

**Incubate and digest protein**

19. Transfer the column to a clean 1.5 mL Lo-bind Eppendorf tube.
20. Add **50 µL** of digestion buffer directly onto the column. **Do not close the tube**. Be careful not to damage the column during pipetting and avoid air bubbles on top of the column.
21. Place the sample in a chamber/box filled with tissue soaked in water to keep the environment inside humid.
22. Incubate samples overnight at 37 °C

**This is the end of day 2**. Please proceed to page 8 to continue the sTRAP protocol to be performed on day 3.

*Roig Kuhn, F., Klaassen, R.V., Koopmans, F.T.W., Koolman, T.S.Z., Smit, A.B., Spijker, S.*
*Dept. of Molecular and Cellular Neurobiology, Vrije Universiteit Amsterdam, Netherlands*

#### Day 3: sTRAP day 2

##### Material:

- Solvent A (0.1% formic Acid in MQ-water)
- Solvent B (0.1% Formic Acid in acetonitrile)
- Speedvac concentrator plus, centrifuge

##### Preparation:

- Check solvents for particles; if needed, refresh the solvent using the stock.

##### Method:

##### Elute and dry peptides

23. Remove the samples from the 37 °C incubator (step 22, Day 2) and inspect the columns to determine whether the digestion buffer has not eluted overnight. If the buffer has eluted, note this observation, as it indicates incomplete protein digestion within the column.
24. Close the tube and centrifuge: 1,450 x g at RT for 1 minute.
25. Add **50 µL** solvent A to the top of each column.
26. Centrifuge column: 1,450 x g at RT for 1 minute.
27. Add **50 µL** solvent B to the top of each column.
28. Centrifuge column: 1,450 x g at RT for 1 minute.
29. Inspect the columns to ensure all the solution has passed through. If any volume remains, repeat centrifugation as needed.
30. Dispose of the column and close the elution tube containing the peptides.
31. Place samples in the speed vac concentrator with the tube caps open. The drying time depends on the number of peptides in the sample; more peptides generally mean a quicker drying time.
32. Setting speed vac: **time:** ∞, **brake:** on, **Temp:** --, **Mode event:** V-AQ
33. If the expected drying time is uncertain, inspect the samples every 2 hours and remove tubes once fully dried. Note down the drying time of each sample.
34. Store the dried peptides at -80 °C.

**This is the end of day 3**<u>. Please proceed to page 9 for the peptide assay to be performed on day 4.</u>

#### Day 4: Peptide Assay

##### Material:

- Stock solutions: 1 M Tris-HCl buffer pH 8.0, HeLa protein digest standard 100 ng/µL (dissolved in solvent A)
- Solvent A (0.1% formic acid in MQ-water)
- 384-well plate
- 1.5 mL Protein Lobind Eppendorf tube
- Centrifuge, thermomixer, microplate scanner

##### Preparation:

- Make a 384-well plate overview
- Check solvent A for particles and refresh if needed.
- Make **2 mL** of 200 mM Tris-HCl buffer pH 8.0

| 200 mM of Tris-HCl buffer pH 8.0 in MQ water | | | |
|---|---|---|---|
| Material | Stock concentration | Required concentration | Volume from stock (mL) |
| Tris-HCl buffer pH 8 (M) | 1 | 0.2 | 0.4 |
| MQ-water (mL) | - | - | 1.6 |
| End Volume | | | 2 |

- Make **20 µL** HeLa standards from the 100 ng/µL HeLa stock:

| HeLa standard (0 - 0.1 - 0.25 - 0.5 - 1.0 µg/µL) | | | | |
|---|---|---|---|---|
| Conc µg/µL | Tris-HCl buffer 200 mM (µL) | Solvent A (µL) | HeLa 100ng stock (µL) | End volume (µL) |
| **0** | 160 | 160.0 | 0.0 | 320 |
| **0.1** | 10 | 9.0 | 1.0 | 20 |
| **0.25** | 10 | 7.5 | 2.5 | 20 |
| **0.5** | 10 | 5.0 | 5.0 | 20 |
| **1** | 10 | 0.0 | 10.0 | 20 |

##### Method:

1. Thaw the dried peptide samples at room temperature.
2. Add solvent A to the dried peptides. The volume of solvent A depends on the initial protein input used for sTRAP. For 10 µg of protein, typically 75 µL of solvent A is added to ensure the peptide concentration falls within the linear range of the HeLa standard curve used in the peptide assay.
3. Place the sample in the thermomixer and mix: 1,800 rpm at RT for 2 minutes.
4. Then switch thermomixer: 2,000 and RT for 2 minutes.
5. Centrifuge sample: 10,000 x g at RT for 1 minute to collect all liquid to the bottom of the tube.
6. Add **20 µL** of 200 mM Tris-HCl buffer pH 8.0, to a labeled 1.5 mL LoBind Eppendorf tube.
7. Add **20 µL** of sample mix to the Tris-HCl buffer; store the remaining peptide mixture (55 µL) in at -80 °C for later use. (Day 5 for Evotips / MS load)
8. Place the **40 µL** sample diluted in Tris-HCl buffer in the thermomixer and mix: 2,000 rpm at RT for 2 minutes.
9. Centrifuge sample: 10,000 x g at RT for 30 seconds to collect all fluid to the bottom of the tube.

10. Place the 384-well plate on ice and pipette **19 µL** HeLa or sample mix into each well. From each sample, measure a **replicate.**
11. Measure the fluorescence at 285 excitation and 355 emission.

#### Settings for the microplate reader:

**Optical configuration:** Monochromator.
**Read mode:** FL
**Read Type**: Endpoint
**Wavelength>** Known> **Excitation:** 9 nm **Emission:** 15 nm
**Number of wavelengths:** 1
**Excitation lm1:** 285 nm **Emission lm1:** 355 nm
**Plate type >**
**Plate format:** 384-wells
**Select specific:** 384-well plate standard opaque

#### Determine the peptide amount in Excel:

#### 1) HeLa standard curve

1. Determine the average absorption for the blanks.
2. Determine the corrected absorption value for each HeLa standard by subtract the average blank from the HeLa standard.

*Absorption value HeLa – average blank = corrected absorption value HeLa*

3. Insert a scatter chart using the values from the corrected absorption of HeLa. Insert a trendline, set the intercept to 0, and display the equation and $R^2$ on the chart

#### 2) Amount of peptide in the sample

4. Determine the average absorption for each sample.
5. Correct the absorption value for each sample.

*Absorption value sample – average blank = corrected absorption value sample*

6. Determine the amount of peptide: **µg/10 µL**.

*Corrected absorption value sample / linear slope = Peptide amount in 10 µL*

7. Determine the amount of peptide: **µg/µL**
8. Convert µg/µL to **ng/µL**

9. Determine the volume (µL) needed from the sample to get 150 ng of peptide for the MS run.

*150 (ng) / concentration (ng/µL) = volume (µL) needed for 150 ng of peptide*

**This is the end of day 4**. Please proceed to page 11 for the Evotip/MS sample loading to be performed on day 5.

*Roig Kuhn, F., Klaassen, R.V., Koopmans, F.T.W., Koolman, T.S.Z., Smit, A.B., Spijker, S.*
*Dept. of Molecular and Cellular Neurobiology, Vrije Universiteit Amsterdam, Netherlands*

#### Day 5: Evotips / MS-sample load

##### Material:

- Solvent A (0.1% formic Acid in MQ-water)
- Solvent B (0.1% Formic Acid in acetonitrile)
- Isopropanol
- Evotips, Evosep box (plus an extra balance Evosep box for the centrifuge)
- 96 well plates and racks for isopropanol & solvent A
- Centrifuge, multi-channel pipette

##### Preparation:

**In this example 3 Evotips are used**

- Calculate the volume amount of sample needed for 150 ng of peptide (see results from peptide assay, step 8 from day 4). Prepare the sample dilution in solvent A if necessary.
- Use 70% ethanol to thoroughly clean the bench surface and remove dust or contaminants
- Inspect all solvent (solvent A, B, and isopropanol) for visible particles that could cause tip blockages. Replace any bottle if contamination is suspected.

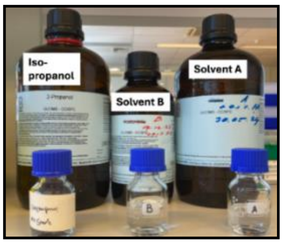


- Place the required number of Evotips in a clean Evosep box. Leave the first two rows empty to allow for proper movement of the tips during sample pipetting.

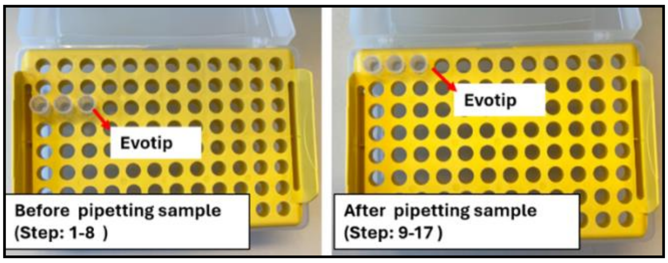


- Prepare a balance Evosep box with the same number of tips to use as a counterweight in the centrifuge.
- Prepare control and reference standards. Make a 50 ng master mix of HeLa standard(from stock: 100 ng/µL, diluted in solvent A) and blanks (solvent a) as negative control.
- With a multi-channel, fill the whole 96-well plate with **150 µl** solvent A per well.
- With a multi-channel, fill the whole 96-well plate with **100 µL** isopropanol per well.
- Set the centrifuge to the correct setting: 800 x g, RT for 1 minute.

*Roig Kuhn, F., Klaassen, R.V., Koopmans, F.T.W., Koolman, T.S.Z., Smit, A.B., Spijker, S.*
*Dept. of Molecular and Cellular Neurobiology, Vrije Universiteit Amsterdam, Netherlands*

#### Method:

I. Thaw the dried peptides mixture at room temperature.
II. Place the sample in the thermomixer and mix: 2,000 rpm at RT for 2 minutes.
III. Centrifuge sample: 10,000 x g at RT for 2 minutes to pellet any potential debris to the bottom of the tube.

**Sample Loading on Evotip**

1. Add **50 µL** of solvent B to each Evotip using a repeating pipette.
2. Centrifuge boxes: 800 x g at RT for 1 minute.
3. Remove the yellow Evotip rack from the Evosep box, empty the waste from the Evosep box and place the rack back into box.
4. Centrifuge again: 800 x g at RT for 1 minute.
   Meanwhile, place the 96-well plate filled with isopropanol ready for step 5.
5. Soak the Evotips in the isopropanol filled 96-well for 10-20 seconds, until the tips turn pale white.
6. Add **50 µL** of solvent A to each Evotip using a repeating pipette.
7. Centrifuge the boxes: 800 x g at RT for 1 minute.
   Meanwhile, replace the isopropanol plate with the 96 well plate containing solvent A for step 8.
8. Transfer the Evotips to the 96-well plate containing solvent A and discard the waste from the Evosep box.

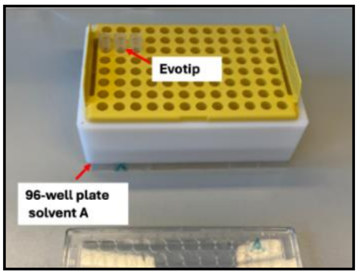


9. Using a standard pipette, add **20 µL** of solvent A into each Evotip. Then, pipette **150 ng** of peptide directly into the solvent.
10. Centrifuge the boxes: 800 x g at RT for 1 minute.

**Wash Evotips (repeat steps 11-13 twice)**

11. Place the Evotip rack into the 96-well plate containing solvent A and discard the waste from the Evosep box.
12. Add **50 µL** of solvent A to each Evotip using the repeating pipette.
13. Centrifuge the boxes: 800 x g at RT for 1 minute.

14. After the second wash, place the Evotip rack into the 96-well plate containing solvent A. Discard waste from the Evosep box.
15. Add **100 µL** of solvent A to the Evotip using a repeating pipette.
16. Centrifuge the boxes: 800 x g at RT for **5 seconds.** (Set centrifuge for 1 minute and manually stop at 55 seconds)
17. Place the Evotip rack back into the 96-well plate containing solvent A and fill the Evosep box halfway with solvent A.
12. Store the Evotips at 4 °C until use or load directly on the MS autosampler.
13. Store the remaining peptide mixture back in the -80 °C freezer.
